# Single-cell profiling reveals pervasive heterogeneity in subcellular RNA localization

**DOI:** 10.64898/2026.02.04.703776

**Authors:** Xiaojuan Fan, Markus Hafner

## Abstract

Subcellular RNA localization is a fundamental layer of gene regulation, yet its heterogeneity across individual cells remains poorly understood. Here, we introduce the RNA Localization Profiler (RLP), a proximity-based RNA-editing strategy that maps compartment-specific RNAs in living cells. Across the cytoplasm, endoplasmic reticulum (ER), and plasma membrane, RLP identifies robust and highly specific RNA localization programs linked to translation and membrane organization. Single-cell RLP (scRLP) reveals that individual cells harbor roughly 5,000-7,000 cytoplasmic RNAs, with <10% associated with the ER. These measurements uncover pervasive subcellular heterogeneity in RNA localization that is undetectable by bulk assays. Spatial RNA patterns define an orthogonal axis of cell-state identity that is independent of gene expression. For example, ZWINT mRNA relocalizes to the cytoplasm in a cell cycle-dependent manner. These findings establish heterogeneous levels of subcellular RNA localization as a variable dimension of intracellular organization and cell identity.

## Introduction

A central question in cell biology is how the crowded intracellular environment is organized in space and time to coordinate complex biochemical reactions. Cells achieve this organization in part by controlling the localization of molecular components: concentrating reactants within specific compartments can accelerate reactions, whereas segregating them can delay or prevent interactions, thereby creating distinct functional microenvironments^1^. RNA localization has emerged as a critical layer of post-transcriptional regulation that fine-tunes gene expression through transcript distribution^2-5^. This process is evolutionarily conserved and essential for development, synaptic plasticity, and memory formation in neurons^6^, and overall rapid adaptation to environmental cues^7,8^. For instance, local translation of mRNAs at cellular protrusions drives polarized motility^9^, a process fundamental to embryogenesis^10^ and cancer metastasis^11,12^. Although transcriptomic studies suggest substantial variability in RNA distribution^13-15^, the extent and nature of this heterogeneity are poorly understood, particularly at single-cell resolution.

Current transcriptome-wide approaches have provided valuable insights but remain constrained by fundamental limitations. Bulk methods such as biochemical or biophysical fractionation^16,17^, fluorescence-activated particle sorting (FAPS)^5^, and proximity labeling^2,18,19^ can identify large sets of compartment-associated RNAs but cannot resolve heterogeneity across thousands of single cells. Imaging-based strategies, including MERFISH^20^ (multiplexed error-robust fluorescence in situ hybridization) and seqFISH+^21^ (sequential FISH), achieve high spatial precision and multiplexing, while TEMPOmap^14^ (temporally resolved *in situ* sequencing and mapping) integrates metabolic labeling with *in situ* sequencing to track RNA dynamics at subcellular resolution. However, these methods require cell fixation, which may alter native RNA distribution, and specialized instrumentation, which limits accessibility. Thus, a scalable strategy to profile RNA localization dynamics in living cells with organelle-level resolution remains an outstanding need.

RNA-editing enzymes have recently been repurposed as molecular recorders to label RNAs in living cells. In TRIBE (targets of RNA binding proteins identified by editing), RNA-binding proteins (RBPs) are fused to the deaminase domain of adenosine deaminase acting on RNA (ADAR) to introduce adenosine-to-inosine (A-to-I) edits on target transcripts^22^. Similarly, the apolipoprotein B mRNA editing catalytic polypeptide 1 (APOBEC1), which catalyzes cytosine-to-uracil (C-to-U) conversion^23^, has been adapted for single-cell mapping of *N*⁶-methyladenosine (scDART-seq, single-cell Deamination Adjacent to RNA Modification Targets Sequencing)^24^ and for antibody-free detection of RBP–RNA interactions (STAMP, Surveying Targets by APOBEC-Mediated Profiling)^25^. More recently, engineered variants of the *Escherichia coli* tRNA-specific adenosine deaminase (TadA) have been applied for RBP–RNA mapping^26^. These studies show that RNA editors can be redirected by fusion partners to leave durable sequence marks on target RNAs, enabling detection by RNA-seq. We reasoned that coupling RNA editors to compartment-specific localization signals could similarly enable scalable, single-cell profiling of RNA localization.

Here, we introduce scRLP, a flexible sequencing-based strategy that quantitatively maps RNA localization in living cells at single-cell resolution. By profiling distinct subcellular compartments and their temporal dynamics, scRLP reveals a previously inaccessible view of how RNA positioning varies across individual cells. These measurements uncover extensive spatial heterogeneity and identify cell states defined by RNA localization rather than gene expression. scRLP therefore establishes a conceptual and quantitative framework for linking subcellular RNA organization to cellular identity and function.

## Results

### Development of the RLP system and validation in the cytoplasm

To track RNA localization dynamics across cellular compartments, we developed the RNA localization profiler (RLP) reporter system. The RLP reporter is built as a single fusion protein comprising three modular components: i) a localization signal that directs the reporter to a specific subcellular compartment; ii) an RNA editing enzyme that marks nearby RNAs through characteristic sequence edits, and iii) an RNA-binding domain (RBD) with high affinity but limited sequence-specificity that promotes efficient reporter-RNA interactions (**Figure 1A**). We expected that in living cells, the RLP would selectively edit RNAs within the targeted compartment for subsequent identification through high-throughput sequencing (RLP-seq). Furthermore, the modular design allows flexible targeting of diverse compartments and is compatible with single-cell analyses.

**Figure 1.**
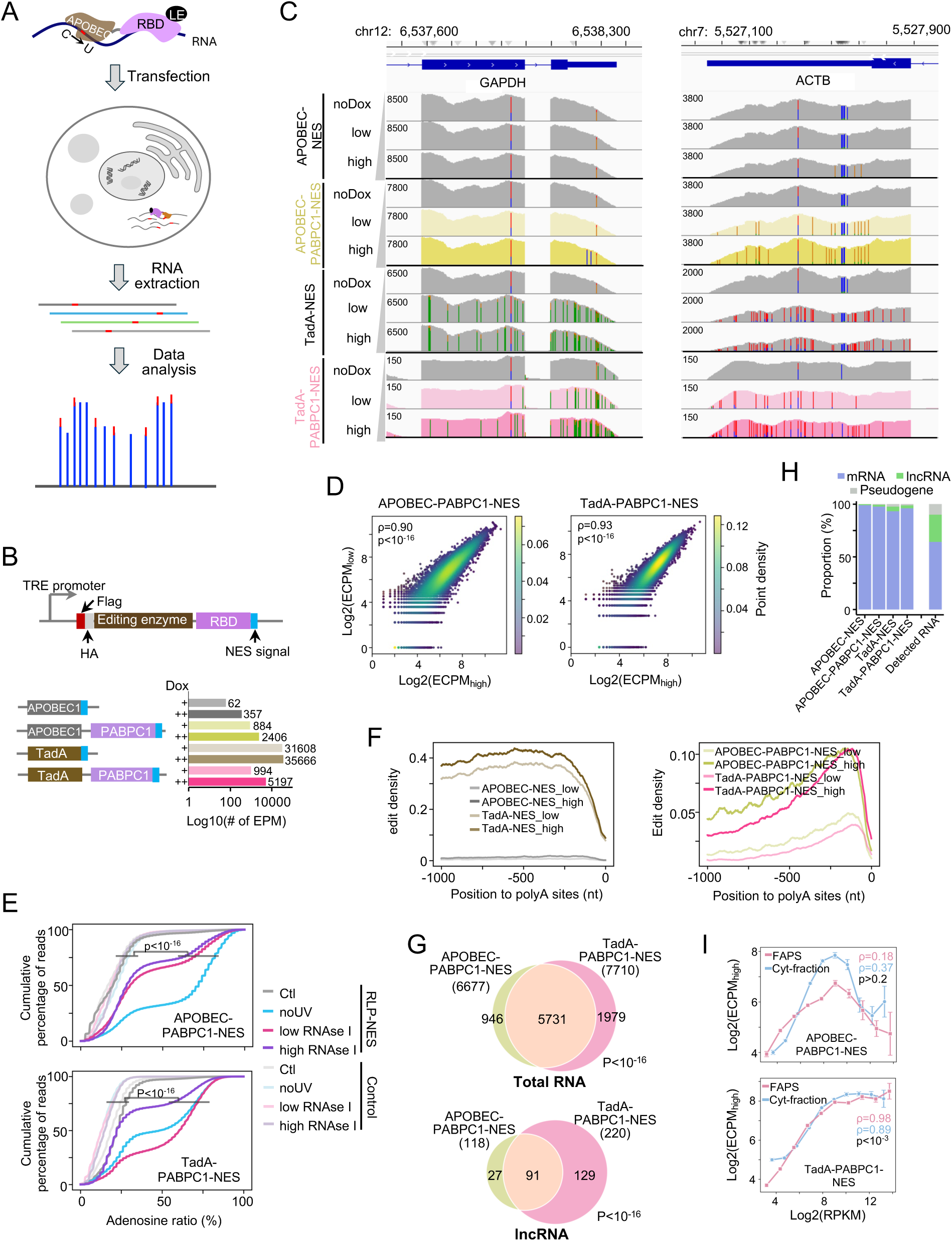
Development of the RLP system and validation in the cytoplasm. **A**, Schematic of the RLP system, comprising a localization element (LE), an RNA-binding domain (RBD), and an RNA editing enzyme, which enables selective tagging of RNAs within targeted subcellular compartments. Edited RNAs are identified by high-throughput sequencing of poly(A)-selected transcripts. **B**, Design of the cytoplasmic RLP reporters using rAPOBEC1 (C-to-U) or TadA8.20 (A-to-I), with or without fusion to the RBD of PABPC1. Control reporters lacking the RBD (APOBEC1-NES, TadA-NES) are shown alongside APOBEC1-PABPC1-NES and TadA-PABPC1-NES. Right, bar plot of editing sites per million mapped reads (EPM, log_10_ scale) under low (+) and high (++) doxycycline induction. **C**, The editing profiles (RNA-seq coverage tracks) of GAPDH and ACTB for APOBEC1-PABPC1-NES (yellow), TadA-PABPC1-NES (pink), and controls (gray) under no, low, or high doxycycline. Colored bars indicate editing events (C-to-U for APOBEC1, A-to-G for TadA; SNV detection threshold = 0.05). Nucleotide colors: A = green, C = blue, G = orange, T = red (reversed on minus strand). **D**, Correlation of gene-level editing abundance (ECPM) between high (x-axis) and low (y-axis) doxycycline induction for APOBEC1-PABPC1-NES (left) and TadA-PABPC1-NES (right). Spearman ρ and p values are indicated. **E**, Cumulative distributions of adenosine content in binding sequences identified by PAR-CLIP for APOBEC1-PABPC1-NES (top) and TadA-PABPC1-NES (bottom). Controls: untransfected cells (gray) and cells transfected with control reporters without UV crosslinking (cyan), with low RNase I (pink), or high RNase I (purple) digestion. Statistical comparisons were performed using the Mann–Whitney U test. **F**, Metagene profiles of editing density relative to poly(A) sites. Left, APOBEC1-NES and TadA-NES controls; right, APOBEC1-PABPC1-NES and TadA-PABPC1-NES. **G**, Overlap of cytoplasmic RNAs detected by APOBEC1-PABPC1-NES and TadA-PABPC1-NES. **H,** Composition of detected cytoplasmic RNAs compared with total RNA-seq. Bars show the proportions of edited mRNAs (blue), lncRNAs (green), and pseudogenes (gray) in APOBEC1-NES, APOBEC1-PABPC1-NES, TadA-NES, TadA-PABPC1-NES, and total detected RNAs by RNA-seq. **I,** Correlation of cytoplasmic RNA abundance measured by APOBEC1-PABPC1-NES/TadA-PABPC1-NES reporters with published cytoplasmic datasets (FAPS^5^ and Cyt-fraction^17^). Spearman ρ and p values are indicated.

We tested two well-characterized RNA editing enzymes in the RLP framework (**Figure 1B; Table S1**): rAPOBEC1, a cytidine deaminase that mediates C-to-U editing^25^, and TadA8.20, a hyperactive variant of the *E. coli* tRNA adenosine deaminase that converts A-to-I on single-stranded RNAs and some structured RNAs^27^. To promote reporter-RNA interactions, the enzymes were fused at their C-termini to the first two RNA recognition motifs (RRMs) of PABPC1, which strongly bind poly(A) tails^28^ present on most mRNAs and many long non-coding RNAs (lncRNAs).

For initial benchmarking, we selected a nuclear export signal (NES) to direct the reporter to the cytoplasm, a compartment for which multiple published datasets obtained using orthogonal methods enabled independent validation^2,5,17^. We used lentiviral integration to generate stable HEK293T cell lines expressing two RLP reporters in a doxycycline-inducible manner, APOBEC1-PABPC1-NES and TadA-PABPC1-NES, along with variants lacking RBDs (APOBEC1-NES and TadA-NES) as controls. Reporter expression and cytoplasmic localization were confirmed using Western blotting and immunofluorescence (IF; **Figures S1A and S1B**). Reporter expression did not affect cell viability and proliferation or the transcriptome profile, except for TadA-NES, which exhibited some cytotoxicity, as evidenced by reduced cell numbers, and greater transcriptomic divergence between uninduced and induced samples (**Figures S1B, S1C**), likely due to extensive cytoplasmic editing activity (**Figures 1B and 1C**). Mutation spectra showed high editing specificity with >98% C-to-U conversions in APOBEC1-PABPC1-NES and nearly 100% A-to-G conversions in TadA-PABPC1-NES (**Figure S1D**). Editing was minimal in uninduced cells (**Figures 1C and S1E**), confirming tight regulation of reporter activity. Finally, RNA-seq detected ∼2,400 editing sites per million mapped reads (EPM) for APOBEC1-PABPC1-NES and ∼5,200 for TadA-PABPC1-NES (**Figure 1B; Table S2**), demonstrating robust activity. Editing output was dose-dependent, with higher doxycycline induction (2 µg/mL) producing more edits and higher rates than low induction (0.1 µg/mL) (**Figures 1B, 1C, and S1E**). Taken together, the RLP system enabled tunable expression and precise control of enzyme dosage and exposure time.

To quantify the abundance of compartment-associated RNAs across conditions, we calculated editing counts per million (ECPM; see Methods), which reflects the proportion of reporter-induced edits per gene and provides a length-independent readout of RNA localization (**Figure S1F**). The editing profiles from APOBEC1-PABPC1-NES and TadA-PABPC1-NES were highly consistent between high- and low-doxycycline induction (Spearman ρ ≥ 0.9, p < 10⁻¹⁶; **Figure 1D**) and significantly correlated between enzymes (Spearman ρ = 0.6, p < 10⁻¹⁶; **Figure S1G**), demonstrating the high reproducibility of the RLP system.

We next asked where edits occurred along transcripts and whether the RBD influenced their position. Compared with APOBEC1-NES, APOBEC1-PABPC1-NES produced >7-fold more edits and >3-fold higher editing density, with strong enrichment in the 3′-UTR and a preference for the expected AC motif at which APOBEC1 is known to edit (**Figures 1B, S1E, and S1H**). In contrast, TadA-PABPC1-NES showed lower activity than TadA-NES but redirected most edits to the 3′-UTR, favoring the expected TA motif (**Figures 1B, S1E, and S1H**). Importantly, inclusion of the PABPC1 RBD prevented cytotoxicity and minimized transcriptional perturbation upon RLP expression (**Figures S1B and S1C**).

We further sought to confirm the RNA-binding specificity of the RLP reporters by PAR-CLIP (photoactivatable ribonucleoside-enhanced crosslinking and immunoprecipitation) to map RLP reporter binding sites at nucleotide resolution^29^ (**Figures S1I and S1J**). We found that RLP-bound sequences were significantly enriched for adenosines relative to controls (cells lacking RLP reporter or reporters without the PABPC1 RRMs), regardless of crosslinking conditions or RNase I digestion intensities (p < 10^-16^; **Figure 1E**). These findings indicate that APOBEC1-PABPC1-NES and TadA-PABPC1-NES preferentially bind polyadenylated RNAs. Consistent with this, editing sites were most frequently located 50–250 nt upstream of the poly(A) tail (**Figure 1F**). Together, these results confirm that the PABPC1-derived RBD drives selective labeling of polyadenylated RNAs near their 3′ ends.

We next defined the cytoplasmic RNAs detected by the RLP system. In total, 6,677 cytoplasmic RNAs were detected by APOBEC1-PABPC1-NES and 7,710 by TadA-PABPC1-NES, with 5,731 shared between the two systems (p < 10^-16^; **Figure 1G; Table S3**). Among these, 118 and 220 lncRNAs were detected by APOBEC1- and TadA-based RLP reporters, respectively, with 91 in common (p<10^-16^; **Figures 1G and S2A**), underscoring the strong concordance between the two RLP systems. Importantly, abundant mitochondrial RNAs or lncRNAs known to be exclusively nuclear were not detected by either reporter (**Table S3; Figure S2B**), indicating the specificity of our system. Transcripts tagged by both APOBEC1 and TadA RLPs had higher expression and accumulated more edits (**Figures S2C and S2D**), suggesting that abundant RNAs are more readily edited. Within this set, TadA-PABPC1-NES consistently produced more edits and higher editing ratios than APOBEC1-PABPC1-NES (**Figures S2D and S2E**), reflecting its stronger enzymatic activity. Overall, cytoplasmic RNAs were >96% mRNAs, <3% lncRNAs, and <1% pseudogenes (**Figure 1H**; **Table S3**), compared to 64%, 26%, and ∼10%, respectively, in total RNA-seq. These results demonstrate that the RLP reporters targeted to the cytoplasm indeed predominantly capture polyadenylated RNA, i.e., mRNAs, as intended (**Figure 1H)**.

To benchmark the RLP system performance in cytoplasm, we compared our data with biochemical Fractionation-seq and FAPS datasets^5,17^ and found comparable numbers and significant overlap of detected RNAs (p < 10^-16^; **Figure S2F**), supporting the system’s ability to comprehensively capture cytoplasmic transcripts. RNA abundance measured by TadA-PABPC1-NES correlated strongly with both cytoplasmic data (Spearman ρ = 0.89-0.98, p < 0.001), whereas the APOBEC1-PABPC1-NES showed only weak correlation (Spearman ρ = 0.16–0.37, n.s.; **Figure 1I**), likely due to its lower intrinsic editing activity (**Figures S2D and S2E**). In the TadA system, the correlation curve plateaued for highly expressed RNAs (log₂RPKM > 10), suggesting that editing saturates at abundantly expressed transcripts (**Figure 1I**). These results indicate that TadA-based RLPs reliably quantify RNA abundance across a broad dynamic range. More broadly, we validated the RLP framework of RNAs by profiling RNAs in the cytoplasm, the compartment with fully orthogonal datasets, and showed that it enables robust, specific, and reproducible identification, establishing a foundation for spatial transcriptome mapping in live cells.

### RLP faithfully captures RNAs at the endoplasmic reticulum membrane

Following cytoplasmic validation, we extended the RLP system to profile RNAs at the endoplasmic reticulum membrane (ERM; **Figure 2A**), a compartment that interfaces with the cytosol and has been extensively characterized^2,5,17,30^. The ERM supports the co-translational synthesis and folding of secretory and membrane proteins and also serves as a global platform for mRNA translation, underscoring its broad role in spatial gene regulation^17^. Conventional approaches typically infer ER localization from relative enrichment over cytosol, which makes it challenging to distinguish true ER-associated RNAs from contamination of abundant cytosolic RNAs. Anchoring an RLP reporter to the cytosolic face of the ER by an ERM signal peptide directly reduces background activity by confining the reporter and integrating editing signals over hours (*vs.* the seconds-scale labeling of APEX), thereby allowing efficient identification of true positive signal from transient encounters.

**Figure 2.**
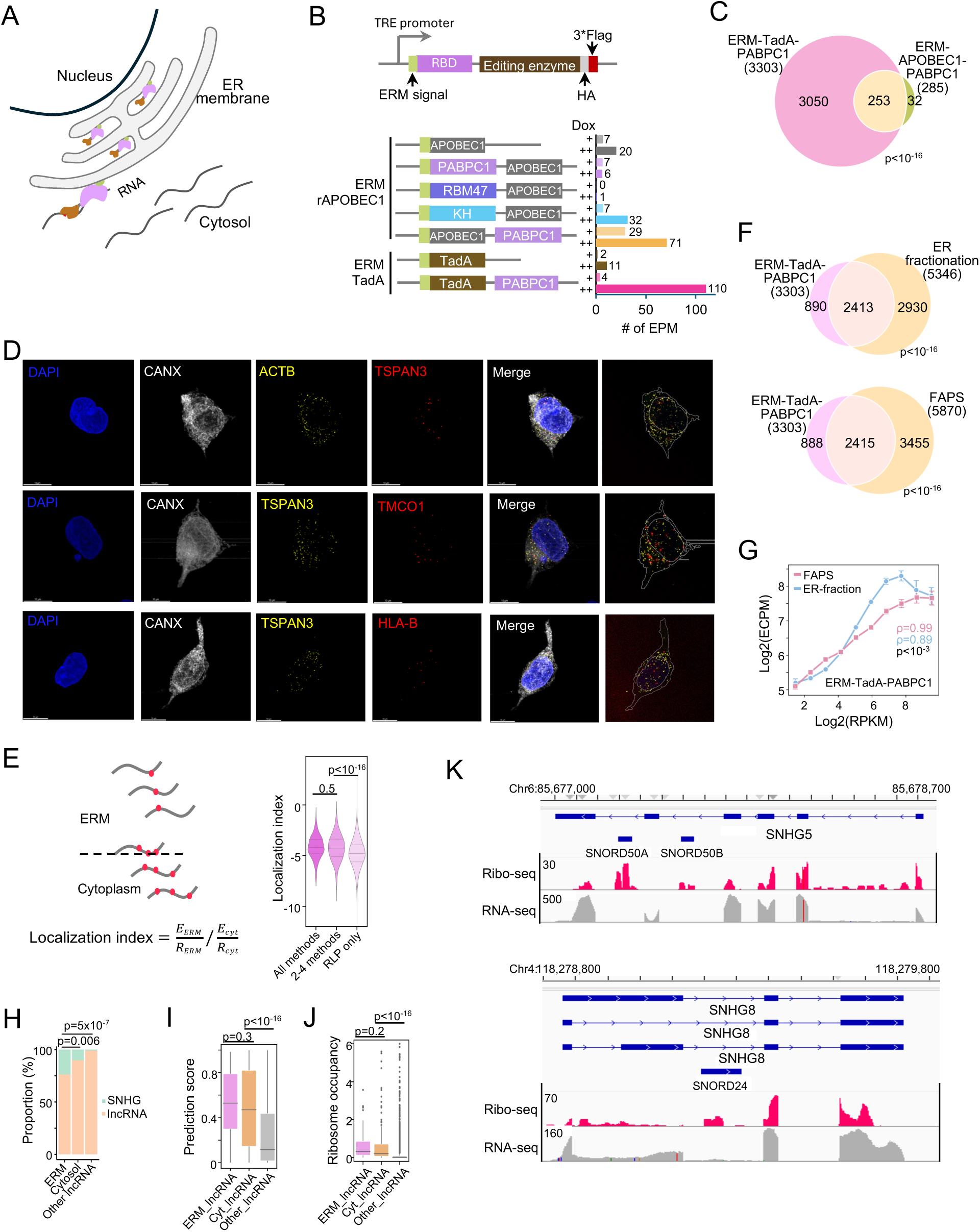
RLP faithfully captures RNAs at the endoplasmic reticulum membrane (ERM). **A**, Schematic of the ERM-targeted RLP reporter. **B**, Design of RLP reporters in the ERM with different RBDs (from PABPC1, RBM47, or KHDRBS3) and positional arrangements fused to APOBEC1 or TadA. Right, bar plot showing EPM under low (+) and high (++) doxycycline induction. **C**, Overlap of RNAs identified by ERM-TadA-PABPC1 and ERM-APOBEC1-PABPC1. **D**, Representative multiplex FISH-IF validation of ERM-associated RNAs from z-stack images. First sample (top row): DAPI (blue), CANX (ER marker, gray), cytosolic control RNA ACTB (yellow), and ERM control RNA TSPAN3 (red). Subsequent samples: DAPI (blue), CANX (gray), ERM control RNA TSPAN3 (yellow), and test ERM RNAs (red; HLA-B, TMCO1). Rightmost panels show nuclear and ER masks (gray) overlaid with RNA puncta (yellow/red). Scale bar, 10 µm. See **Supplementary Videos 1–3**. **E**, Schematic of the localization index (LI), defined as the ratio of edit counts (normalized by read density) in ERM *vs.* cytoplasm (left). Right, distribution of LI for RNAs detected by all five methods (n=158), 2-4 methods (n=587), or uniquely by ERM-TadA-PABPC1 (RLP-only, n=2,558). **F**, Overlap of RNAs identified by ERM-TadA-PABPC1 with published ER RNA datasets^5,17^ under relaxed criteria (no ER/cytosol enrichment cutoff). **G**, Correlation between ER RNA abundance measured by ERM-TadA-PABPC1 and published ER datasets^5,17^. Spearman ρ and p value are indicated. **H**, Fraction of SNHG family lncRNAs detected by ERM-TadA-PABPC1 (ERM_lncRNA, n=62), TadA-PABPC1-NES (Cytosol_lncRNA, n=247), and other lncRNAs (n=19,640). P values were determined by Fisher’s exact test. **I-J**, TranslationAI-predicted translation initiation scores (panel **I**) and ribosome occupancy (panel **J**) for lncRNAs detected by ERM-TadA-PABPC1, TadA-PABPC1-NES, and other lncRNAs. **K**, RNA-seq (gray) and Ribo-seq (red) coverage tracks for SNHG5 and SNHG8.

To optimize the RLP reporter, we again varied the RBD, its placement within the construct, and editing enzyme (**Figure 2B**). In addition to the previously validated RRM domains from PABPC1 to recruit the RLP to mRNA, we also tested the KH domain of KHDRBS3 and the RRM domains of RBM47, each of which recognizes the AAUAAA motif^31^, the most abundant hexamer in the human transcriptome (**Table S1)**. The different constructs showed widely varying editing activity (**Figure 2B**), confirming that PABPC1 RRMs positioned at the C termini of the reporters provide the most effective recruitment for both APOBEC1- and TadA-based reporters (**Figure 2B**). Notably, all ERM-targeted reporters with corresponding cytoplasmic counterparts (ERM–APOBEC1–PABPC1, ERM–TadA–PABPC1, and their RBD-deficient counterparts) exhibited substantially lower editing activity, showing ∼17- to 3,000-fold reductions in EPM (**Figures 1B and 2B**). Nearly all (>98%) ERM-localized RNAs were also detected in the cytoplasm, including 282 of 285 from ERM–APOBEC1–PABPC1 and all 3,303 from ERM–TadA–PABPC1, but with significantly fewer editing events and lower editing ratios (**Figures S3A and S3B**). These reductions likely reflected the confinement of the reporter to the ER membrane to yield lower overall output but more spatially specific editing at the ERM.

Among ERM constructs, ERM-TadA-PABPC1 showed the highest editing activity (>100 EPM, **Figure 2B**; **Table S2**), with ∼90% RNAs identified by ERM-APOBEC1-PABPC1 also recovered by ERM-TadA-PABPC1 (**Figure 2C**). Therefore, ERM-TadA-PABPC1 was selected for subsequent experiments. Western blotting and IF confirmed proper expression and localization of the ERM-TadA-PABPC1 reporter at the ERM (**Figures S3C and S3D**). At the transcript level, edits were enriched near transcript 3′-ends (**Figure S3E**), consistent with the poly(A)-tail binding specificity of PABPC1. Sequencing of ERM-TadA-PABPC1 detected 3,303 ERM-localized RNAs, including 3,217 protein-coding RNAs (>97%), 62 lncRNAs (∼2%), and 24 pseudogenes (<1%) (**Figures 2C and S3F; Table S2**). Localization of representative transcripts (e.g., HLA-B, TMCO1, SNHG8, and NSA2) was further validated by FISH-IF imaging (**Figures 2D and S3G; Supplementary Videos 1-5**).

The RLP-identified RNAs showed significant overlap with APEX-seq, ER-fractionation, FAPS, and MERFISH datasets (p < 10^-16^, hypergeometric test; **Figure S3H**). To quantify compartment enrichment at the transcript level, we defined a localization index (LI) as the ratio of edit counts, normalized by read density, in the ERM relative to the cytoplasm (**Figure 2E**, see Methods). Transcripts detected at the ERM by multiple independent methods showed significantly higher LI than those uniquely identified by ERM-TadA-PABPC1 (**Figure 2E**), supporting the LI as a robust measure of compartment-specific RNA localization.

Unlike these snapshot approaches, ERM-TadA-PABPC1 integrates editing over 24 hours, providing a cumulative record of stably ER-associated RNAs. To benchmark ERM-TadA-PABPC1, we reanalyzed ER fractionation^17^ and FAPS^5^ datasets without applying ER/cytosol enrichment cutoffs. Under the relaxed criteria, ∼2,400 RNAs overlapped with the ∼3,300 RNAs detected by ERM-TadA-PABPC1 (5,346 by ER fractionation and 5,870 by FAPS; **Figure 2F**). This highly significant overlap (p < 10^-16^) indicates that ERM-TadA-PABPC1 captures *bona fide* ERM-localized RNAs, including many excluded by strict enrichment thresholds. RNA abundance measured by ERM-TadA-PABPC1 correlated strongly with ER fractionation as well as FAPS (Spearman ρ = 0.89–0.99, p < 0.001; **Figure 2G**), collectively supporting the accuracy of all the methods, considering that they are based on completely independent principles of RNA isolation. Nevertheless, RLP showed higher specificity, as e.g., abundant cytosolic markers such as GAPDH and ACTB were absent in RLP data (**Figure S3I**), whereas traditional methods (Fractionation-seq or FAPS) assigned ∼20% of their signal to the ER fraction. Similarly, abundant mitochondrial RNAs were also not present in our data, emphasizing the ability of RLP to accurately define the ERM-localized transcriptome.

As the ER membrane is known to translate mRNAs coding for secretory, membrane-associated, and some cytosolic proteins, we did not expect to find many lncRNAs in this compartment. Nevertheless, 62 lncRNAs were significantly edited by ERM-TadA-PABPC1, including 15 from the small nucleolar RNA host gene (SNHG) family (**Figures 2H and S3J**). Additionally, the ERM-localized SNHGs showed stronger editing than other ERM-localized lncRNAs (**Figure S3K**), prompting us to test their translational potential at the ER. TranslationAI, a deep-learning model for translation site prediction^32^, assigned significantly higher initiation scores to ERM- and cytoplasm-localized lncRNAs than to other lncRNAs (**Figure 2I**). These predictions were corroborated by ribosome profiling (**Figures 2J and 2K**). Together, these findings align with recent evidence for the translational potential of SNHGs^33^ and reveal a previously unappreciated subset of translationally active transcripts misannotated as lncRNAs at the ER membrane and in the cytosol, and further underscore the ability of the RLP system to faithfully map RNA localization.

### Time-course analysis of ERM-localized RNA dynamics

After confirming the specificity of the RLP system at the ER membrane, we next investigated RNA localization dynamics at this compartment over a time course of 72 h. ERM-TadA-PABPC1 expression peaked at 12 h and declined thereafter, while editing reached maximal density at 24 h and remained 3′-biased (**Figures 3A, 3B, S4A, and S4D, Table S2**). Although reporter expression was detectable by 2 h, editing remained negligible (**Figures 3A and S4A–S4D**), indicating a slowly accumulating detectable editing signal, which minimized consideration of spurious signals from brief encounters. Across all time points, 4,421 ERM-associated RNAs were identified, of which 3,237 were reproducibly observed (**Figure 3C, Table S4**). Comparison with the cytoplasmic pool (∼8,600 RNAs) revealed that nearly half of the cytoplasmic transcripts associate with the ER, either consistently or intermittently, indicating widespread ER association at the bulk-cell level.

**Figure 3.**
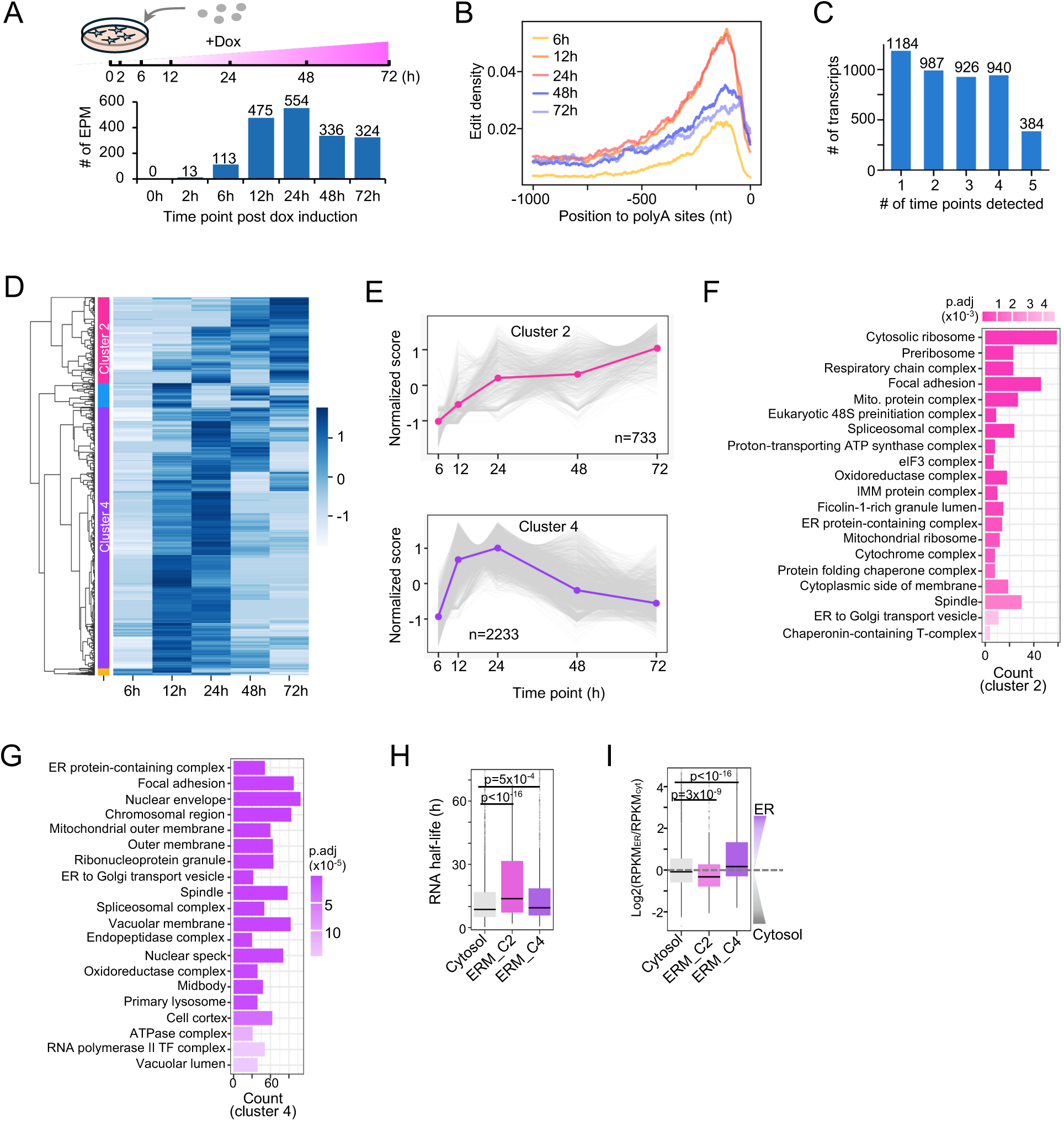
Time-course analysis of ERM-localized RNA dynamics. **A**, Experimental design (top) and bar plot of EPM from 0–72 h after ERM-TadA-PABPC1 induction (bottom). **B**, Distribution of edit density across 1,000 nt upstream of poly(A) sites at the indicated time points (6–72 h). **C**, Number of RNAs reproducibly detected across different numbers of time points. **D**, Heatmap of Z-score–normalized LI for ERM-associated RNAs across the ERM-TadA-PABPC1 time course. Rows represent genes; columns represent time points. Hierarchical clustering groups RNAs by temporal dynamics. Cluster identities are indicated by the left color bar (C2, pink; C4, purple). **E**, Localization dynamics for C2 and C4 RNAs. Gray traces, individual genes; colored lines, cluster means. **F-G**, GO Cellular Component enrichment for C2 (panel **F**) and C4 (panel **G**). Bar color indicates adjusted p values. Abbreviations: Mito., mitochondria; IMM, inner mitochondrial membrane. **H**, RNA half-lives for cytosolic (not detected in ERM), C2, and C4 RNAs. **I**, RNA enrichment scores (log_2_[RPKM-ER/RPKM-cytosol]) from a published ER/cytosol fractionation dataset.

To resolve temporal patterns, we clustered localization index (LI) values across time points (**Figure 3D, Table S4**), revealing two major clusters with distinct kinetics and functions. Cluster 2 (C2, 733 RNAs) showed steadily increasing ERM association from 6 to 72 hours (**Figures 3E, 3F, and S4E**) and was enriched for translation and mitochondrial pathways (**Figure 3F**). Cluster 4 (C4, 2,233 RNAs) mirrored ERM-TadA-PABPC1 editing dynamics (**Figures 3A, 3E, 3F and S5A**) and was linked to endomembrane organization and transport (**Figure 3G**). Notably, cytosolic markers GAPDH and ACTB showed no ERM-associated edits throughout the time course (**Figure S5B**), demonstrating that the RLP system distinguishes ERM-associated RNAs from abundant cytosolic transcripts with high specificity.

The distinct dynamics of C2 and C4 likely arise from two factors: RNA turnover and reporter distribution. Although C2 RNAs generally have longer half-lives, most RNAs are short-lived (**Figure 3H**), suggesting that reporter localization dynamics primarily drive the clustering patterns. Consistently, particle sorting data^5^ showed that C2 RNAs were also found in the cytosol to considerable levels, whereas C4 RNAs preferentially localized to the ER (**Figure 3I**). Despite their presence in the cytosol, C2 RNAs showed rising LI, dense 3′-biased edits, and FISH validation, confirming their dynamic interaction with the ER. Thus, ERM-TadA-PABPC1 resolves both stable and intermittent RNA–ER interactions, capturing a broad spectrum of ER-associated transcript dynamics with high specificity.

### Application of the RLP system to the plasma membrane

With the scalability and specificity of the RLP system established, we next applied the system to the plasma membrane (PM), a compartment inaccessible to conventional fractionation but proposed to harbor localized translation and RNA-mediated signaling that regulate processes such as cell adhesion, migration, and intercellular communication^9,11,34^. As an anchor, we selected the gap junction protein GJA1 (connexin-43), a well-characterized membrane protein that mediates direct cell–cell communication and small-molecule exchange through gap junction channels and hemichannels. GJA1 undergoes a defined trafficking cycle—synthesized in the ER, oligomerized in the Golgi, and delivered to the PM—where its turnover is tightly regulated by ubiquitination, clathrin-mediated endocytosis, and either lysosomal degradation or recycling via AMSH-dependent deubiquitination^35^ (**Figure 4A**). Importantly, both C- and N-termini of GJA1 face the cytosol, allowing the editing module of the RLP reporter to be fused at either end to label RNAs at the cytosolic surface of the PM. Combined with its spatial restriction at cell–cell contact sites and central role in communication and signaling, these features make GJA1 an ideal anchor for dissecting RNA dynamics at the PM using the RLP system.

**Figure 4.**
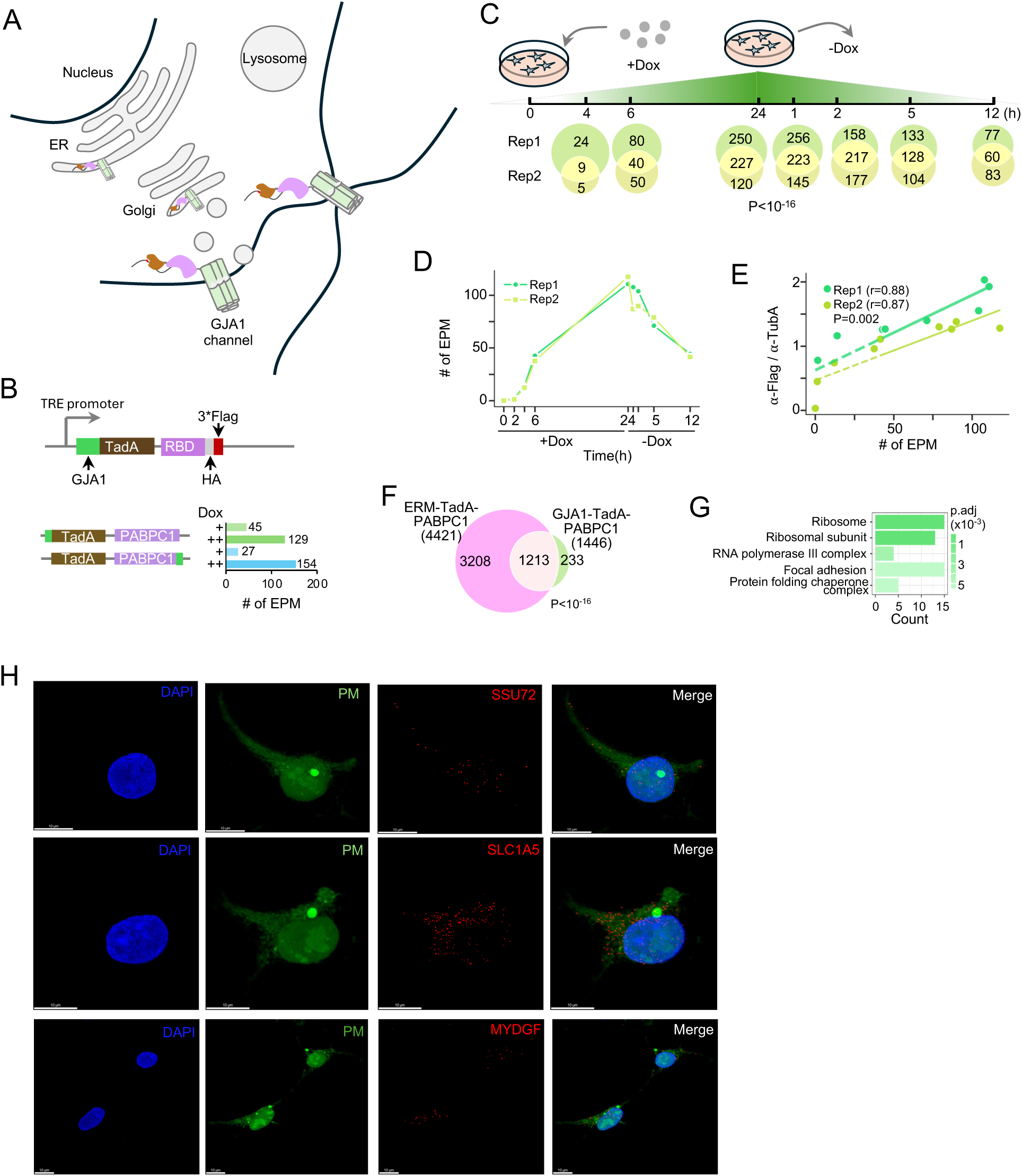
Application of the RLP system to the plasma membrane (PM). **A**, Schematic of the GJA1 (connexin 43) life cycle, from ER synthesis and Golgi trafficking to PM assembly, turnover, and recycling. **B**, Design of PM-targeted GJA1-TadA-PABPC1 reporters in which TadA-PABPC1 was fused to GJA1 at the N terminus (GJA1–TadA–PABPC1) or C terminus (TadA–PABPC1– GJA1). Right, bar plot of the EPM under low (+) or high (++) doxycycline induction. **C**, Dual-phase time-course experiment. Top, schematic of induction (0–24h) and withdrawal (1–12h) phases. Bottom, overlap of edited RNAs between biological replicates. **D**, EPM across the dual-phase time course in two replicates. **E**, Correlation between editing sites and GJA1-TadA-PABPC1 expression (normalized to α-TUBA) across time points. Linear fits are shown; Pearson ρ=0.88 (Rep1) and ρ=0.87 (Rep2); p=0.002. **F**, Overlap of RNAs detected by GJA1-TadA-PABPC1 and ERM-TadA-PABPC1 across all time points. **G**, GO Cellular Component enrichment for PM-localized (GJA1-TadA-PABPC1-specific) RNAs. **H**, Multiplex FISH validation of PM-associated RNAs from z-stack images. DAPI (blue), entire cell (green), and test RNAs (red; SSU72 and MYDGF). Scale bar, 10 µm. See **Supplementary Videos 7-9**.

Building on the superior performance of TadA-based RLP reporters over rAPOBEC1 in both cytoplasmic and ERM compartments (**Figures 1–2**), we engineered stable HEK293 cell lines inducibly expressing GJA1–TadA–PABPC1 (GJA1 fused to the N terminus) or TadA–PABPC1–GJA1 (GJA1 fused to the C terminus) to target cell–cell interfaces (**Figure 4B; Table S1**). Both constructs efficiently edited RNA (>100 EPM, **Figure 4B; Table S2**), identifying 598 and 568 RNAs, respectively, with 285 shared transcripts (∼50% overlap; p < 10⁻¹⁶; **Figure S6A**). Nevertheless, the GJA1-TadA-PABPC1 reporter showed slightly higher editing efficiency with a tighter 3′-end bias of the editing signature (**Figure S6B**), and was selected for subsequent analyses.

To capture the localization dynamics of RNAs associated with the PM, we performed a time-course experiment in biological duplicates where we harvested cells at 0, 2, 4, 6, and 24 h during doxycycline induction to allow reporter accumulation, and at 1, 2, 5, and 12 h after withdrawal to halt new synthesis (**Figures 4C** and **S6C**). This strategy was designed to monitor reporter trafficking to the membrane and identify time points with enriched reporter abundance at the PM. As expected, GJA1-TadA-PABPC1 protein levels increased steadily during induction and declined rapidly after doxycycline removal (**Figures S6C-S6F**), with most reporters localizing from cytoplasm to the PM by 1 h after withdrawal, consistent with the rapid turnover rate of GJA1^36^ (**Figure S6F; Supplementary Video 6**). The editing activity closely tracked reporter abundance across all time points in both replicates (π > 0.87, p < 0.001; **Figures 4D and 4E; Table S2**). Despite temporal fluctuations in expression and localization, transcript sets identified at each time point showed highly significant overlap between replicates (p < 10^-16^; **Figure 4C**), underscoring the stability of PM-associated RNA organization and the robustness of the GJA1-TadA-PABPC1 in capturing dynamic localization patterns.

Given that GJA1 is synthesized at the ER before trafficking to the PM, we expected substantial overlap between RNAs tagged by GJA1–TadA–PABPC1 and ERM–TadA–PABPC1. Indeed, the two datasets largely coincided (>80%; **Figure 4F**). The remaining ∼16% (233 RNAs) were unique to GJA1-TadA-PABPC1, with 60 (>25%) corresponding to genes whose protein products have been detected by membrane pulldown assays^4^, suggesting localized translation at the cell surface. These unique RNAs were enriched for Gene Ontology terms related to protein synthesis, transcription regulation, focal adhesion, and protein folding (**Figures 4G, Table S5**), consistent with the functional categories of proteins identified in the same dataset^4^. Representative RNAs uniquely captured by GJA1-TadA-PABPC1, including SSU72, SLC1A5, and MYDGF, were further validated by FISH (**Figure 4H, Supplementary Videos 7-9**).

### Characteristics of RNAs in distinct compartments

With the set of compartment-specific RNAs in hand, we compared their sequence-specific properties. We first examined translational features of compartment-specific RNAs, and analyzed the codon adaptation index (CAI), a proxy for codon optimality linked to translation efficiency and RNA stability. Among ERM-associated transcripts, RNAs with an editing profile that mirrored reporter levels (cluster 4, C4, **Figure 3**) exhibited significantly lower CAI values than RNAs stably associated with the ERM (cluster 2, C2, **Figure 3**) or cytosolic RNAs (**Figure 5A**), yet showed higher ribosome occupancy (**Figure 5B**). This inverse relationship may reflect specialized co-translational control: C4 RNAs are enriched for transmembrane domains and signal peptides (**Figure 5C**) and may use suboptimal codons to slow elongation, enabling proper co-translational folding and membrane insertion^37^. Supporting this role, C4 RNAs also possessed the longest 3′- UTRs (**Figure 5E**), consistent with their role as *cis*-acting ER localization signals^38^. Evolutionary analysis further showed that C4 RNAs were the most conserved set (**Figure 5D**), indicating selective pressure to maintain these specialized features. By contrast, across all analyses, including CAI, ribosome occupancy, transmembrane enrichment, and conservation, C2 RNAs more closely resembled cytosolic transcripts, suggesting they represent an intermediate population bridging cytosolic and ER-associated RNAs.

**Figure 5.**
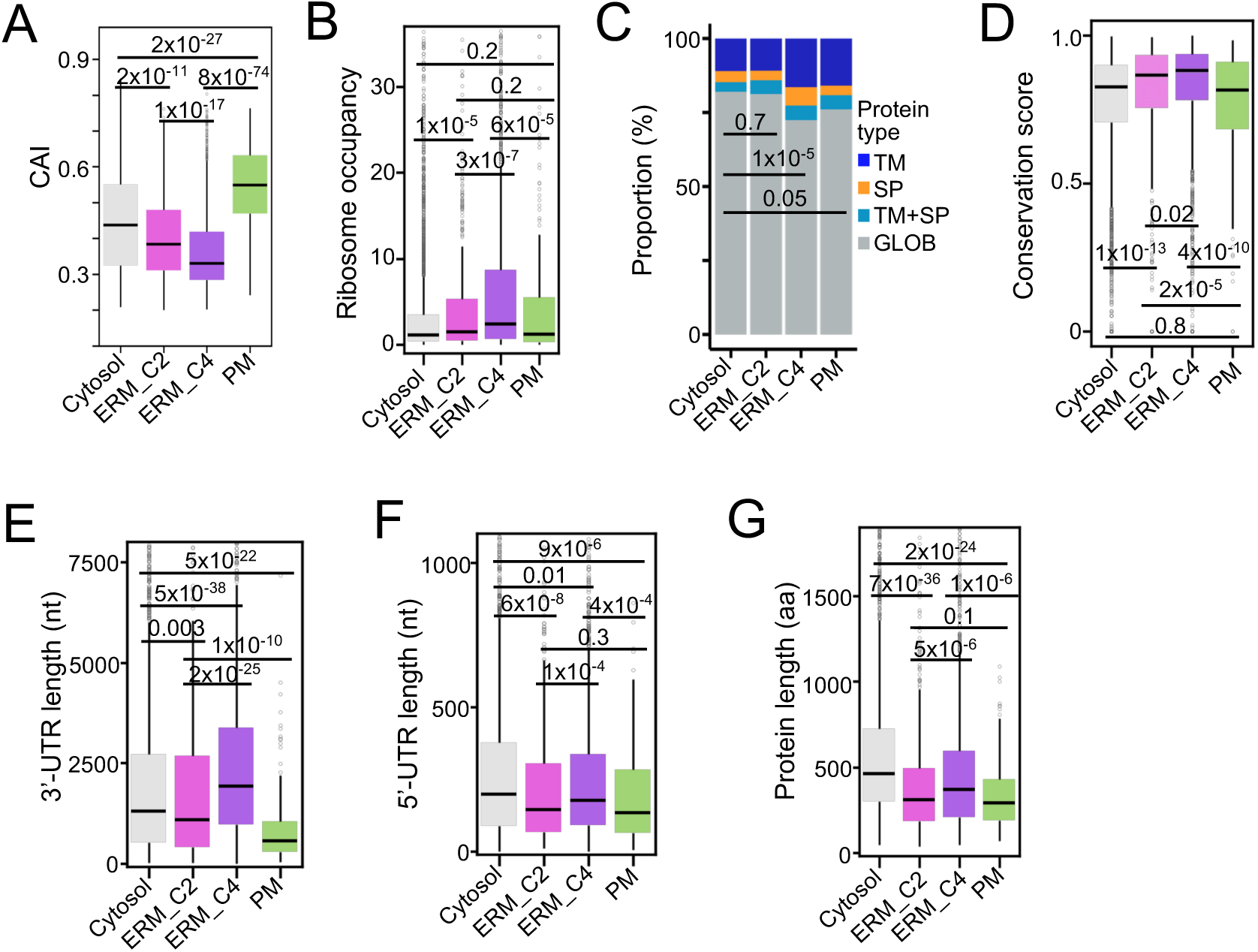
Distinct molecular features of cytosolic, ERM, and PM-localized RNAs. Comparison of **A**, codon adaptation index (CAI), **B**, ribosome occupancy, **C**, proportion of predicted domains, **D**, conservation score, **E**, 3′-UTR length, **F**, 5′-UTR length, and **G**, protein length across cytosolic (n=4,002), C2 (n=733), C4 (n=2,233), and PM RNAs (n=233). Predicted protein features were classified by DeepTMHMM as transmembrane domains (TM), signal peptides (SP), or globular proteins (GLOB).

PM–associated RNAs identified by GJA1-TadA-PABPC1 displayed the highest CAI values (**Figure 5A**), suggesting intrinsic optimization for rapid translation. They also had the shortest 5′-UTRs, coding sequences, and 3′-UTRs (**Figures 5E-G**), features previously linked to increased ribosome density on shorter ORFs^39^. Consistently, their ribosome occupancy was ∼5-fold higher than that of cytosolic RNAs (**Figure 5B**), suggesting that these compact, codon-optimized transcripts are tuned for rapid translation at the dynamic plasma membrane. Together, these findings show that RNAs in distinct subcellular compartments carry unique molecular signatures reflecting their specialized functions, underscoring the precision of RLP system in defining compartment-specific localization.

### scRLP-seq identifies cytoplasmic RNAs in single cells

The ultimate objective driving the development of the RLP system was the resolution of RNA localization at the single-cell level (single-cell RLP sequencing, scRLP-seq). First, we performed scRLP in stable cells expressing TadA-PABPC1-NES. Using SMART-seq2^40^, which provides high sensitivity for comprehensive transcript detection, we profiled 979 single cells (∼5 million paired-end reads per cell; **Figure 6A**). Across cells, ∼90% of single-nucleotide variants corresponded to TadA-mediated A-to-G conversions (**Figure S7A**), confirming high editing specificity. Edits were enriched within 200 nt upstream of transcript 3′-ends, independent of local read depth (**Figure 6B**). Gene-level read densities from pseudobulk scRLP-seq closely matched those from bulk RLP-seq (Spearman ρ = 0.88, p < 10⁻¹⁶; **Figure S7B**), and per-gene ECPM values were also significantly correlated (Spearman ρ = 0.77, p < 10⁻¹⁶; **Figure S7C**). These results demonstrate that scRLP-seq faithfully recapitulates bulk RLP-seq profiles, confirming robust reproducibility across modalities.

**Figure 6.**
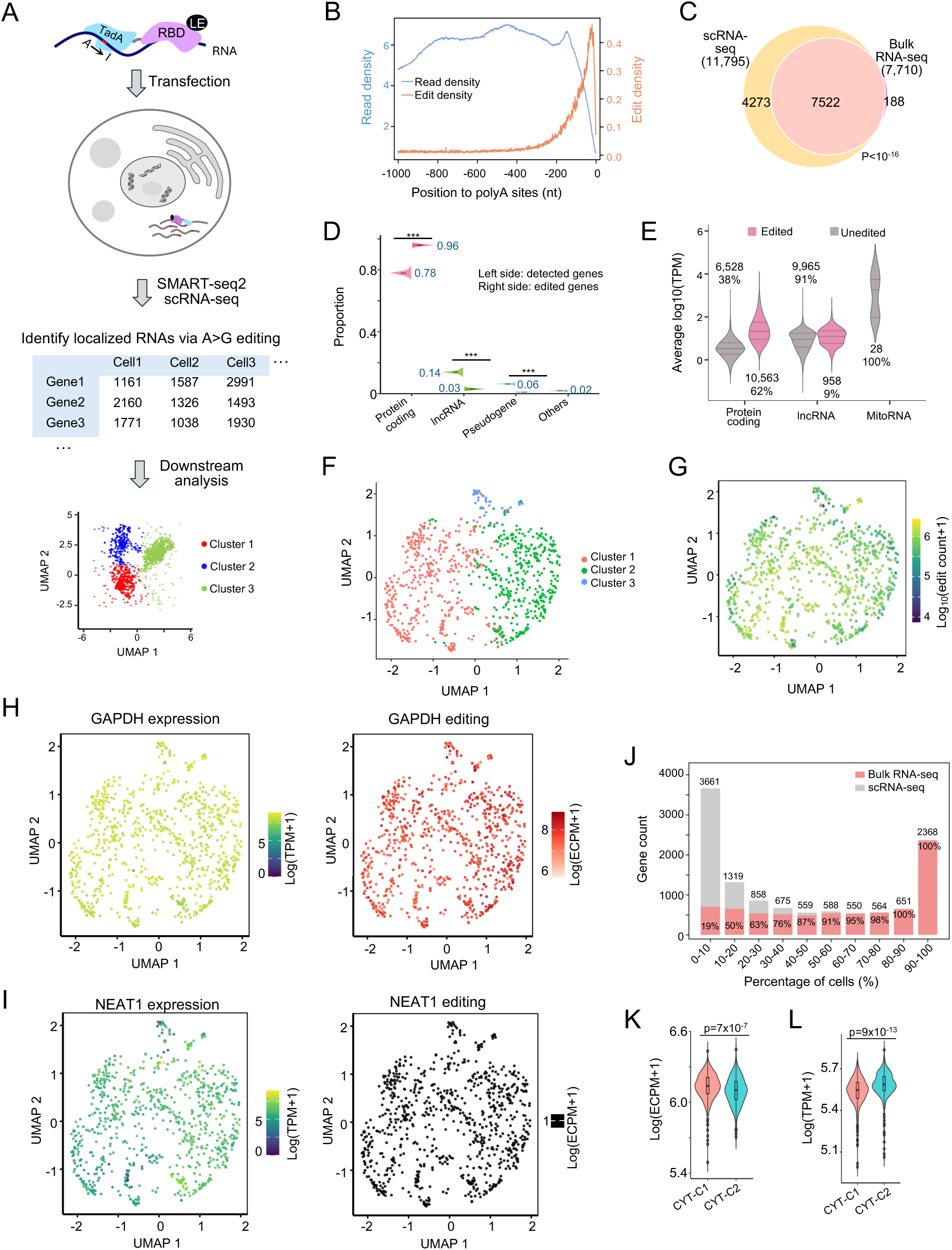
scRLP-seq identifies cytoplasmic RNAs in single cells. **A**, Workflow schematic of scRLP-seq using the TadA-PABPC1-NES reporter with SMART-seq2 for full-length transcript profiling. **B**, Read density (left, blue) and edit density (right, orange) within 1,000 nt upstream of poly(A) sites. **C**, Overlap of cytoplasmic RNAs detected by scRLP-seq and bulk RLP-seq. **D**, RNA biotype distributions across single cells, comparing detected RNAs (left) with edited RNAs (right). Categories: protein-coding, lncRNA, pseudogene, other. **E**, Expression distributions of protein coding, lncRNAs, and mitochondrial RNAs, separated into edited (pink) and unedited (gray) transcripts. **F**, UMAP of single cells based on transcriptome-wide expression. **G**, Same UMAP as in panel **F**, overlaid with per-cell editing levels. **H–I**, UMAPs from panel **F** overlaid with expression and editing of GAPDH (panel **H**), or NEAT1 (panel **I**). **J**, Edited gene counts (y-axis) binned by detection frequency across cells (x-axis). Gray, scRLP-seq; pink, overlap with bulk RLP-seq. Total gene counts (top) and overlap percentages (bottom) are indicated. **K-L**, Editing levels (panel **K**) and expression (panel **L**) of the top 200 most variable genes (from **Figure S7J**) across CYT-C1 and CYT-C2 clusters.

Across 806 high-quality cells (see Methods for filtering parameters), we identified 11,795 cytoplasmic RNAs, with >97% of bulk-detected RNAs recovered by scRLP-seq (**Figure 6C, Table S6**). The number of edits and edited genes per cell scaled with sequencing depth (Spearman ρ > 0.6, p < 10^-16^, **Figures S7D and S7E**), indicating that detection remained within a broad dynamic range rather than approaching saturation. On average, ∼5,000 cytoplasmic RNAs were detected per cell, with up to ∼7,000 in some cells (**Figure S7E**). This number is comparable to the ∼8,000 cytoplasmic RNAs identified in bulk RLP-seq, suggesting that cytoplasmic RNAs detected in bulk are broadly distributed across individual cells rather than restricted to specific subpopulations. Edited RNAs were strongly enriched for protein-coding genes, comprising ∼96% of edited genes compared with ∼78% in the total detected RNA set from the same scRLP-seq dataset (**Figure 6D**). By contrast, abundant mitochondrial RNAs showed no detectable editing (**Figure 6E**), indicating that subcellular localization, rather than expression level, primarily determines editing. Consistently, editing was independent of expression-based clustering (**Figures 6F and 6G**): the cytoplasmic marker GAPDH was robustly edited in all cells (**Figure 6H**), whereas the nuclear lncRNA NEAT1 showed none (**Figure 6I**). Together, these results demonstrate that coupling RLP with SMART-seq2 enables robust and reproducible mapping of RNA localization at single-cell resolution.

scRLP-seq enabled us to quantify cell-to-cell heterogeneity in cytoplasmic RNA localization. Across 11,795 detected cytoplasmic RNAs, we observed a U-shaped distribution: 2,368 RNAs (20%) were detected in >90% of cells, whereas 3,661 RNAs (31%) appeared in <10% (**Figure 6J**). This pattern closely mirrored that of ∼36,000 RNAs detected by scRNA-seq (**Figure S7F**), which profiles total cellular transcripts including nuclear RNAs. The similarity indicates that cytoplasmic RNA localization heterogeneity parallels overall transcript variability across cells, reflecting true biological variation rather than technical bias. Chi-square testing against expression-matched random simulations confirmed significant deviation from the null hypothesis (p < 10^-16^; **Figure S7G**), indicating localization-specific rather than abundance-driven RNA targeting. RNAs edited in >90% of cells were all recovered in bulk RLP-seq, whereas <20% of those edited in <10% of cells overlapped with the bulk data (**Figure 6J**), underscoring the sensitivity of scRLP-seq. Functionally, broadly edited RNAs (>90% of cells) were enriched for essential processes such as gene expression, RNA processing, mitochondrial metabolism, and protein folding (**Figure S7H**). By contrast, rarely edited RNAs (<10% of cells) were enriched for developmental and morphogenetic pathways, such as morphogenesis, neural development, cell migration, and signal transduction (**Figure S7I**). Rare editing was mainly driven by low expression, but localization-specific regulation also contributed, as many highly expressed RNAs remained rarely edited (**Figure S7J**).

To further examine cell-to-cell variation in cytoplasmic RNAs, we performed UMAP analysis of editing profiles, which separated cells into two clusters (CYT-C1 and CYT-C2; **Figure S7K**). Both expression- and editing-based clustering highlighted genes involved in RNA processing and metabolism (**Table S7)**. Notably, CYT-C1 showed higher editing levels despite lower overall expression than CYT-C2 (**Figures 6K and 6L),** indicating that cytoplasmic RNA distribution can vary independently of transcript abundance.

To investigate RNA localization dynamics across cellular states, we profiled cytoplasmic RNAs in single cells stratified into G1, S, and G2/M phases using marker gene expression. Across the cell cycle, 232 RNAs (significantly enriched for cell cycle processes) showed significant changes in editing levels that correlated strongly with gene expression (Spearman ρ = 0.97, p < 10⁻¹⁶; **Figure S8A**), indicating that most observed RNA changes likely arise from upstream transcriptional regulation rather than altered localization.

However, ZWINT mRNA diverged from this pattern: its cytoplasmic editing increased markedly during G2/M, despite no change in overall abundance (**Figures S8B and S8C**). ZWINT encodes a core component of the outer kinetochore KNL1 complex, which anchors spindle assembly checkpoint proteins and mediates microtubule–kinetochore attachments^41^. During interphase, kinetochores are confined to the nucleus but very briefly become cytoplasm-accessible following nuclear envelope breakdown at prophase. Thus, the increased cytoplasmic editing of ZWINT during G2/M likely reflects a regulated nuclear-to-cytoplasmic redistribution of its mRNA, rather than undetectable passive leakage during nuclear envelope breakdown. Because intron detention is a typical mechanism for nuclear mRNA retention and has been proposed to create a reservoir of transcripts that can be rapidly released upon cellular cues^42,43^, we next tested whether ZWINT localization is controlled by its introns. We quantified intron retention across single cells in S and G2/M phases using an intron ratio (see Methods) for each constitutive intron. Notably, in S phase, introns 2 and 3 showed >10-fold higher ratios than other introns in polyadenylated ZWINT mRNA, indicating strong nuclear detention (**Figure S8D**). These detained introns decreased significantly in G2/M (p < 10^-16^; **Figure S8D**), supporting a model in which post-transcriptional splicing enables cytoplasmic export of mature ZWINT mRNA in this phase. Consistently, introns 2 and 3 were previously identified as periodically spliced retained introns during the cell cycle in bulk RNA-seq analysis^44^. Furthermore, APEX2-mediated proximity labeling of nuclear domain–associated RNAs^45^ revealed that introns 2 and 3 of ZWINT were enriched in nuclear fractions, particularly in SRSF1-targeted nuclear speckles (**Figures S8E and S8F**), which retain intron-containing transcripts differentially regulated across diverse cellular contexts, including the cell cycle^45^. Together, these findings suggest that ZWINT mRNA is likely retained in the nucleus during interphase and redistributed to the cytoplasm in G2/M through regulated splicing of its detained introns, exemplifying how RNA relocalization may coordinate functional timing independently of new transcription.

### scRLP-seq identifies ERM associated RNAs in single cells

To profile ERM-localized transcripts at single-cell resolution, we performed scRLP-seq on HEK293T cells stably expressing inducible ERM-TadA-PABPC1. We profiled 1,406 single cells at an average depth of ∼8 million paired end reads per cell. Across cells, gene-level read densities from pseudobulk scRLP-seq were strongly correlated with bulk RLP-seq (Spearman ρ = 0.89, p < 10⁻¹⁶; **Figure S9A**) and editing density per gene also correlated between pseudobulk and bulk datasets (Spearman ρ = 0.53, p < 10⁻¹⁶; **Figure S9B**), supporting cross-platform consistency. From 713 high-quality cells, we identified 6,568 ERM-associated RNAs, recovering 90% of those detected in bulk RLP-seq (**Figure 7A; Table S8**).

**Figure 7.**
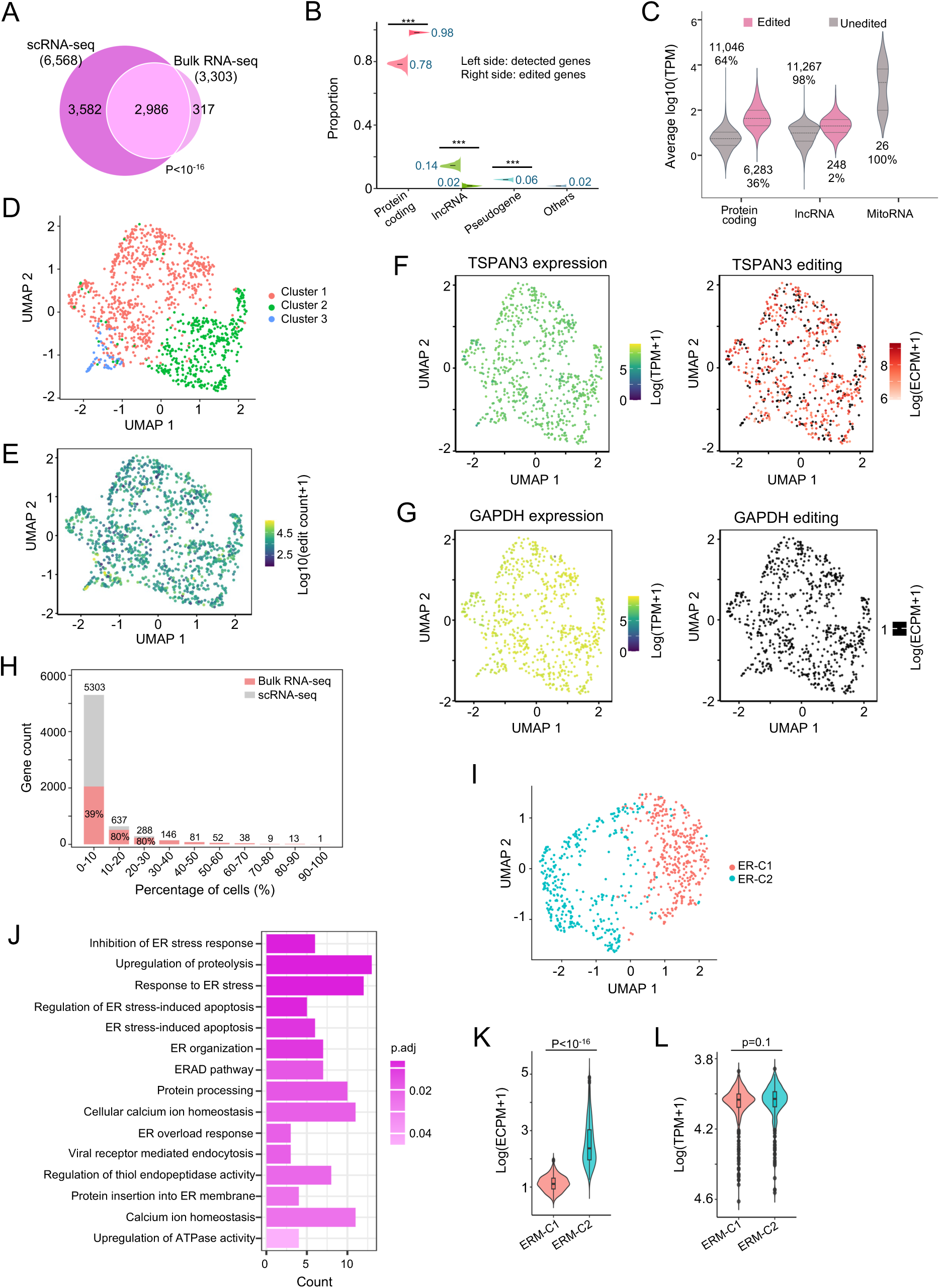
scRLP-seq identifies ERM-associated RNAs in single cells. **A**, Overlap of ERM-associated RNAs detected by scRLP-seq and bulk RLP-seq (ERM-TadA-PABPC1). **B**, RNA biotype distributions across single cells, comparing detected RNAs (left) with edited RNAs (right). Categories: protein-coding, lncRNA, pseudogene, other. **C**, Expression distribution of protein coding, lncRNAs, and mitochondrial RNAs, separated into edited (pink) and unedited (gray) transcripts. **D**, UMAP of single cells based on transcriptome-wide expression. **E**, Same UMAP as in panel **D**, overlaid with per-cell editing levels. **F–G**, UMAPs from panel **D** overlaid with expression and editing of TSPAN3 (panel **F**) or GAPDH (panel **G**). **H**, Edited gene counts (y-axis) binned by detection frequency across cells (x-axis). Gray, scRLP-seq; pink, overlap with bulk RLP-seq. Total gene counts (top) and overlap percentages (bottom) are indicated. **I**, UMAP of single cells based on editing profiles. **J**, GO analysis of the top 200 most variable genes from the editing-based UMAP in panel **I**. Bar color indicates statistical significance (p values). **K-L**, Editing levels (panel **K**) and expression (panel **L**) of the top 200 most variable genes from panel **I** across ERM-C1 and ERM-C2 clusters.

Unexpectedly, the total number of edits and edited genes per cell correlated only weakly with sequencing depth (Spearman ρ < 0.2; **Figures S9C and S9D**), suggesting that ERM-associated editing was largely saturated under our conditions. On average, ∼450 ERM-associated RNAs were detected per cell, with up to ∼1,400 in individual cells (**Figure S9D**). This corresponds to <10% of the ∼5,000 cytoplasmic RNAs per cell, despite bulk RLP-seq showing that ∼50% of cytoplasmic transcripts contact the ERM at some point. Such heterogeneity cannot be explained by clonal variation, as all cells used in scRLP-seq originated from a single FACS-sorted progenitor clone.

When comparing gene categories, the proportion of edited protein-coding genes in scRLP-ERM (98%) was even higher than in scRLP-NES (96%) (**Figures 6D and 7B**), indicating an enrichment of translated RNAs at the ER. As with scRLP in cytoplasm, abundant mitochondrial RNAs remained unedited (**Figure 7C**) and RNA expression levels across cells did not correlate with editing status (**Figures 7D and 7E**). For instance, the ERM marker TSPAN3 was edited in 67% of single cells (**Figure 7F**), whereas the highly expressed cytosolic marker GAPDH showed none (**Figure 7G**). Together, these findings underscore the specificity of scRLP-seq in resolving compartment-associated RNAs at single-cell resolution.

To assess heterogeneity across cells, we quantified the distribution of ERM-associated RNAs within the population. Surprisingly, >80% of RNAs were detected in the ERM of fewer than 10% of cells, whereas only ∼5% appeared in more than 30% (**Figure 7H**). This heterogeneity is unlikely due to sequencing bias in RNA detection, as the occurrence distribution of all RNAs detected by scRNA-seq was comparable between ERM and cytoplasmic regions (**Figures S7F** and **S9F**), indicating that it reflects scRLP-specific editing rather than scRNA-seq bias. Similar distributions were observed across cell-cycle phases (**Figure S9E**), indicating that this heterogeneity is consistent across subpopulations. Chi-square testing against expression-matched simulations confirmed significant deviation from the null (p < 10^-16^; **Figure S9G**), supporting localization-specific rather than abundance-driven targeting. Additionally, ∼40% of RNAs present in the ERM of <10% of cells were also edited in bulk RLP-seq (**Figure 7H**), suggesting that many ERM-associated transcripts are highly dynamic.

To dissect cell-to-cell variation in ERM-associated RNA localization, we performed UMAP analysis of editing profiles, which separated cells into two clusters, ERM-C1 and ERM-C2 (**Figure 7I**). Unlike expression-based clustering, where the top variable genes were enriched for cell cycle processes (**Figure S9H**), editing-based clustering highlighted genes involved in ER stress responses (**Figure 7J)**. Editing levels differed significantly between ERM-C1 and ERM-C2 (p < 10⁻¹⁶; **Figure 7K**), whereas gene expression showed no significant differences (**Figure 7L),** revealing heterogeneity in ER-associated RNA regulation independent of transcript abundance. More broadly, this highlights how single-cell RNA localization profiling can expose functional cell states that conventional expression analyses fail to resolve.

## Discussion

The scRLP system provides a robust framework for mapping RNA localization in living cells with temporal and single-cell resolution. Its reproducibility is supported by strong correlation of editing signatures across RLP reporter levels, concordance between bulk and pseudobulk data, consistent overlap across time-course replicates, and agreement with orthogonal datasets. Its specificity arises from the intrinsic slower kinetics of RNA editing, which minimize detections of transient and spurious contacts compared with seconds-scale APEX labeling. Consistently, abundant cytosolic RNAs (e.g., GAPDH and ACTB) were absent from ERM profiles, and mitochondrial RNAs were undetected in both cytoplasm and ERM profiles, even after extended labeling or at single-cell resolution. RLP scalability and accuracy are demonstrated by the detection of ∼8,600 cytoplasmic RNAs–a number one would expect to be expressed in cell lines-broad concordance with conventional approaches, and the discovery of translationally active (misannotated) lncRNAs such as SNHGs in the ER and cytoplasm. Finally, RLP is flexible: we successfully targeted the cytoplasm, ERM, and PM, and the same design can be extended to other compartments and adapted to diverse experimental conditions, cell types, and tissues. Together, these features establish scRLP as a versatile and standardized platform for dissecting RNA localization dynamics in living cells.

Single-cell RLP profiling revealed striking differences between bulk and single-cell views of RNA organization. Whereas bulk RLP-seq data suggest widespread transcript expression in the cytoplasm and broad ERM association, single-cell measurements uncovered extensive heterogeneity, with only a small core of RNAs consistently localized at the same place across all assayed cells and many others detected sporadically. This divergence highlights the limitations of population-averaged approaches and demonstrates that RNA localization is far more dynamic and variable than previously appreciated. Importantly, localization patterns alone were sufficient to stratify cells into distinct subpopulations, independent of their expression, uncovering spatially defined states that are invisible to conventional transcriptomics. The cell-cycle-regulated relocalization of ZWINT mRNA exemplifies this principle, illustrating how RNA redistribution may be involved in the temporal control of cellular processes without changes in transcript abundance.

Beyond resolving single-cell heterogeneity, the scRLP system provides a framework to dissect the spatiotemporal dynamics and functional principles of RNA localization. Unlike conventional methods that capture static snapshots, scRLP integrates editing events over time, enabling dynamic processes to be resolved in living cells with minimal perturbation. Our analyses show that RNAs exhibit compartment- and cluster-specific signatures ranging from codon usage and ribosome occupancy to RNA stability and evolutionary conservation, which may align with their spatial roles. This selective organization underscores how RNA localization contributes to localized translation, trafficking, and signaling. More broadly, scRLP maps the transcriptome with organelle-level precision in single cells, uncovering a previously hidden layer of spatial organization that shapes cellular states and provides a simple and easy-to-use platform for probing fundamental principles of RNA compartmentalization.

Despite these strengths, several limitations remain. Because the RLP reporter relies on recombinant expression of fusion constructs, direct application to primary tissues remains challenging. Stringent criteria for calling editing events may underestimate the localization of low-abundance transcripts or weakly edited sites, and sequencing depth directly affects the sensitivity of editing detection. In addition, because RNA editing proceeds with relatively slow kinetics, transient or low-affinity reporter–RNA interactions are likely to be missed, thereby limiting the capture of short-lived localization events. Accurate identification of compartment-specific RNAs further depends on the use of appropriate controls to remove unintended labeling; for example, PM-associated RNAs must be distinguished from transcripts encountered during GJA1-TadA-PABPC1 synthesis and trafficking. This caveat applies to all reporter-based approaches for mapping RNA localization unless reporter synthesis or turnover can be selectively inhibited.

In summary, the RLP system opens new opportunities for probing spatial RNA regulation in living cells at single-cell resolution. Coupling RLP with perturbation screens, live-cell imaging, or multi-omics will extend its utility and provide mechanistic insights into how RNA localization contributes to cell identity, developmental programs, stress adaptation, and disease progression. As additional compartments and cell types are profiled, the method can be refined to generate increasingly precise and comprehensive localization maps. Together, these advances position RLP as a foundation for exploring the spatial dimension of gene regulation.

## Methods

### Plasmid constructs and transfection

All cloning was performed according to the enzyme manufacturer’s protocols unless otherwise noted. The plasmid pLIX403_Capture1_APOBEC_HA_P2A_mRuby (a gift from Eugene Yeo; Addgene #183901)^25^ served as the starting template. To avoid potential interference with reporter localization caused by the P2A and mRuby sequences, we generated plasmid pLIX403_Capture1_APOBEC1_HA by PCR-based removal of these sequences using Q5 DNA polymerase (NEB).

#### Control and cytoplasmic reporters

For APOBEC1-NES, the NES signal was fused to the C terminus of APOBEC1, and an N-terminal Flag-HA tag was introduced by Q5 site-directed mutagenesis (NEB). For RLP-NES reporters, first two RRMs from human PABPC1 (ORFeome clone OHS5894-202497981) linked by an XTEN spacer were PCR-amplified and inserted downstream of APOBEC1 in the Flag_HA_APOBEC1_NES backbone using the ClonExpress Ultra One Step Cloning Kit (Vazyme). TadA-based reporters (Flag_HA_TadA_NES and Flag_HA_TadA_PABPC1_NES) were created by replacing APOBEC1 with TadA8.20, amplified from pET28a-6xHis-MBP-TEV-TadA8.20 (a gift from Weixin Tang; Addgene #194702)^27^.

#### ERM-targeted reporters

RBDs (PABPC1, KHDRBS3, or RBM47) were fused either upstream or downstream of APOBEC1. For upstream RBD constructs (ERM-RBD-APOBEC1-HA-3xFlag), a 3×Flag tag was first introduced at the C terminus of APOBEC1_HA, and synthesized ERM signal with RBDs were inserted upstream using Gateway BP/LR cloning (Invitrogen). For downstream constructs (ERM-APOBEC1-PABPC1-HA-3xFlag), the ERM signal was fused to APOBEC1_HA_3xFlag by Gateway cloning, and XTEN-linked PABPC1 RRMs were inserted downstream by one-step cloning. TadA-based ERM reporters were generated in the same manner.

#### Plasma membrane reporters

Two PM-targeted reporters were generated by fusing TadA-PABPC1 to full-length human GJA1 (a gift from Alice Ting, Addgene plasmid # 49385)^46^. In one design (GJA1-TadA-PABPC1-HA-3xFlag), GJA1 was transferred via Gateway recombination into APOBEC1_HA_3xFlag. In a second (Flag_HA_TadA_PABPC1_GJA1), GJA1 was inserted downstream of TadA_PABPC1 using a Gateway attR1-ccdB-attR2 cassette. All primers and synthesized sequences are listed in **Table S1**.

### Cell culture and generation of stable cell lines

All stable RLP cell lines were generated using human HEK293T cells (Invitrogen), which are derived from transformed female human embryonic kidney tissue. Cells were cultured in Dulbecco’s Modified Eagle Medium (DMEM; Gibco) supplemented with 10% tetracycline negative fetal bovine serum and 100 U/ml penicillin, and 100 μg/ml streptomycin at 37 °C with 5% CO_2_. Cells were passaged at 70–90% confluency by dissociating with TrypLE Express Enzyme (Gibco).

#### Lentivirus production

For each 10-cm dish at ∼70% confluence, HEK293T cells were transfected with 1.64 pmol reporter plasmid, 1.3 pmol psPAX2 (a gift from Didier Trono, Addgene #12260), and 0.72 pmol pMD2.G (a gift from Didier Trono, Addgene #12259) using PEI (Polysciences, DNA:PEI ratio 1:2). Viral supernatants were harvested at 48 and 72 h post-transfection, pooled, and filtered through 0.45-µm PVDF filters (Millipore).

#### Stable cell lines

HEK293T cells were seeded at ∼50% confluency and transduced with 200, 400, 600, and 800 µL viral supernatant supplemented with polybrene (Sigma) at a final concentration of 10 µg/mL. After 24 h, medium was refreshed, and after 48 h, puromycin (Thermo Fisher, 2 µg/mL) was applied until stable populations were established. To further enrich for cells expressing the desired RLP or control reporters and avoid heterogeneity introduced by population-level transfection variation, single cells were isolated via fluorescence-activated cell sorting (FACS), clonally expanded, and validated for subsequent experiments.

### Fluorescence-activated cell sorting (FACS)

Single-cell sorting was performed on a BD FACSAria III cell sorter (BD Biosciences, serial number P656700000039) operated with BD FACSDiva software (v9.0.1). Stable HEK293T reporter populations were harvested at ∼70% confluence, dissociated with TrypLE Express (Gibco), and resuspended in ice-cold PBS. Sorting gates were defined using forward and side scatter (FSC/SSC) parameters to select single, viable cells. DAPI was added immediately prior to sorting to exclude dead cells, and live cells were identified as DAPI⁻ events. Single viable cells were sorted in single-cell mode into 96-well plates containing pre-warmed DMEM for clonal expansion.

### Induction of RLP reporters and RNA editing assays

Stable HEK293T cell lines expressing RLP reporters (APOBEC1-PABPC1-NES, TadA-PABPC1-NES, ERM-TadA-PABPC1, GJA1-TadA-PABPC1, APOBEC1-NES, and TadA-NES) were induced with doxycycline (Sigma) at 0.1 µg/mL (low dose) or 2 µg/mL (high dose) for 24 h. RNA was extracted using TRIzol (Thermo Fisher) and purified with the Direct-zol RNA Miniprep Kit (Zymo Research). Matched uninduced controls were processed in parallel.

Time-course experiments. For ERM-TadA-PABPC1, stable cells were induced with doxycycline (2 µg/mL) and harvested at 0, 2, 6, 12, 24, 48, and 72 h. For GJA1-TadA-PABPC1, cells were induced (2 µg/mL doxycycline) and collected at 0, 2, 4, 6, and 24 h. Induction was terminated by twice PBS wash and fresh DMEM medium replacement, and cells were harvested at 1, 2, 5, and 12 h post-washout. Short post-withdrawal intervals were selected to account for the rapid turnover of GJA1^36^.

### Immunoblotting

Cells were lysed in RIPA buffer (50 mM Tris pH 8.0, 150 mM NaCl, 1% NP-40, 0.5% sodium deoxycholate, 0.1% SDS, 5 mM EDTA) for 10 min on ice, and lysates were cleared by centrifugation (15,000 g, 10 min, 4 °C). Proteins were resolved on NuPAGE 4–12% Bis-Tris gels (Thermo Fisher) and transferred to PVDF membranes. Primary antibodies: anti-Flag (Sigma F1804, 1:2000), anti-TUBA4A (Sigma T9026, 1:2000), anti-GAPDH (ProteinTech 10494-1-AP, 1:20,000). Secondary HRP-conjugated antibodies (Thermo Fisher) were used at 1:5000. Signal intensities were quantified in Fiji.

### Immunofluorescence staining

Stable HEK293T cells were seeded in 8-well chamber slides (ibidi) at 5×10^4^ cells/mL (300 µL/well) and induced with doxycycline (2 µg/mL) prior to fixation. For standard immunofluorescence, cells were treated for 24 h prior to fixation. For time-course analysis, doxycycline was added 72, 48, 24, 12, 6, 2, or 0 h before fixation, so that all samples were fixed simultaneously. Cells were rinsed with PBS, fixed in freshly prepared 2% paraformaldehyde (10 min, RT), permeabilized/blocked in PBS with 0.04% saponin, 5% BSA, and 1% normal goat serum (1 h, RT), and incubated overnight at 4 °C with primary antibodies diluted in blocking buffer. After three washes in PBS with 0.04% saponin, cells were incubated with Alexa Fluor–conjugated secondary antibodies (Invitrogen; 1:500 in blocking buffer, 2 h, RT, dark). Nuclei were counterstained with Hoechst (1:200) during the second wash. The stained cells were kept in 1× PBS prior to imaging.

Primary antibody dilutions: Anti-Flag antibody (Sigma, #F1804) diluted 1:1500 (for APOBEC1-PABPC1-NES and TadA-PABPC1-NES) or 1:500 (for ERM-TadA-PABPC1 and GJA1-TadA-PABPC1 reporters). Anti-CANX antibody (Thermo Fisher, #PA5-34754) diluted 1:500. Phalloidin–Alexa Fluor 647 conjugate (Invitrogen, #A22287) diluted 1:500 was used for F-actin staining.

### RNA fluorescence in situ hybridization (RNA FISH)

RNAs were visualized using the QuantiGene ViewRNA ISH Cell Assay (Thermo Fisher) according to the manufacturer’s protocol with modifications. Cells were seeded on poly-L-lysine–coated glass-bottom chambered dishes (ibidi) at 70–80% confluence, fixed in freshly prepared 4% paraformaldehyde for 30 min at RT, permeabilized with Detergent Solution QC for 5 min, and treated with Protease QS (1:4,000 dilution) for 10 min at RT. Probe hybridization was performed at 40 °C for 3 h using custom probe sets, followed by sequential incubations with PreAmplifier, Amplifier, and Label Probe mixes (each 30 min at 40 °C) with intervening washes.

For ERM-localized RNAs, samples were post-fixed in 4% paraformaldehyde for 15 min at RT, blocked in blocking buffer (3% BSA with 0.1% Tween-20 in PBS) for 30 min at RT, incubated with anti-CANX (Thermo Fisher, 1:500) for 1 h at RT in blocking buffer. After washing, cells were incubated with Alexa Fluor–conjugated secondary antibody (Goat anti-Rabbit IgG (H+L), Alexa Fluor 488, Invitrogen, 1:500) in the same blocking buffer for 30 min at RT, counterstained with DAPI, washed, and stored in PBS at 4 °C protected from light until imaging.

For PM-localized RNAs, nuclei were counterstained with DAPI (1:100 dilution) for 1 min. Entire cells were labeled with HCS CellMask Deep Red (1:50,000, Thermo Fisher) for 10 min. Cells were stored in PBS at 4 °C protected from light until imaging.

### Microscopy

Confocal super-resolution microscopy was performed using a Leica TCS X SP8 microscope driven by the Leica LAS X software and equipped with a 405 nm solid state laser for DAPI and a Leica White Light Laser (470-670 nm) for fluorophores emitting in the Vis spectrum. Acquisitions were performed using either a HC Plan-Apochromat CS2 40X/1.3NA or a 63X/1.4NA oil immersion objective (Leica) through the LIGHTNING mode to achieve sub-diffraction resolution. Single plane micrographs (1,024 pixel wide) were acquired in frame mode to avoid crosstalk among fluorescence channels, with line averaging set at 3-4, and pinhole at 0.4-0.5 Airy units. Z-stacks acquisitions were performed over an axial range set *ad hoc* to cover the whole thickness of the imaged cells with Z steps set at no more than 0.4μm. Images and stacks were saved as .lof files and then exported as .tiff with lossless compression. Linear adjustments to individual channel intensities, when needed, and 3D renderings or maximum intensity projections from Z stacks were performed either on the Leica LAS X or on the Imaris 10.2 (Bitplane) software.

To quantify colocalization between CANX (green) and FLAG (red) signals, 3D Z-stacks acquired on the Leica TCS SP8 were analyzed using the Imaris Colocalization module. The entire cell volume was defined as the region of interest (ROI). Automatic thresholding was applied to remove background fluorescence. Pearson’s correlation coefficient was calculated across all voxels within the ROI to assess the linear correlation between the two channels.

### PAR-CLIP experiments and analysis

PAR-CLIP was performed following the procedure described previously^47^, with minor modifications detailed below. Stable HEK293T cell lines expressing APOBEC1-PABPC1-NES, TadA-PABPC1-NES, or control reporters (APOBEC1-NES and TadA-NES) were used; untransfected controls were included. Cells were induced by adding 2 µg/mL doxycycline for 24 hours and labeled with 4-thiouridine (100 μM) 16 h before UV crosslinking at 365 nm with 0.5 J/cm^2^ (CL-1000 Ultraviolet Crosslinker). Cells were subsequently lysed on ice in 3 mL IP buffer (20 mM Tris pH 7.5, 150 mM NaCl, 2 mM EDTA, 1% NP-40, 0.5mM DTT, and protease inhibitors cocktail (1 tablet/10ml)) for 10 min followed by centrifugation at 15,000 × g for 15 min. Cleared lysates were incubated with anti-FLAG M2 magnetic beads (Sigma, Cat# M8823) for 2 hours at 4°C to immunoprecipitate APOBEC1-PABPC1-NES, TadA-PABPC1-NES, and control reporters. After IP, the beads were treated with RNase I (Thermo Scientific) at low (0.015 U/µL) or high (0.15 U/µL) concentrations for 10 minutes. Next, RNAs bound to the beads were ligated directly to a fluorescently labeled, pre-adenylated 3′ RNA adaptor using T4 RNA Ligase 2, truncated K227Q (NEB) overnight at 4°C.

For SDS-PAGE purification, immunoprecipitated proteins were denatured in 2× loading buffer and heated at 95°C. Supernatants were separated on LDS-PAGE gels alongside diluted PageRuler Plus markers. After electrophoresis, gels were scanned for AF647, aligned with printed images, and bands corresponding to the protein plus ∼25 kDa were excised. Gel slices were recovered using gel breaker tubes and centrifugation, with corresponding regions from untransfected controls processed in parallel. The 3′ adaptor-ligated RNA footprints were eluted by Proteinase K digestion and the released footprints were then ligated to a 5′ RNA adaptor using T4 RNA Ligase 1 (Thermo Fisher). Adaptor-ligated RNA was reverse-transcribed into cDNA using SuperScript IV reverse transcriptase (Invitrogen). The resulting cDNA libraries were PCR-amplified, purified, and quantified prior to sequencing. Libraries were sequenced on an Illumina NextSeq 2000 platform with single-end 50-nucleotide reads. Base calling and conversion to FASTQ format were performed using the Illumina bcl2fastq (v2.20.0) software.

As RLP-NES reporters use the PABPC1-derived RNA-binding domain that specifically recognizes poly(A) tails, captured sequences often contain polyadenylated regions that can’t align to the human genome. To address this limitation, we quantified adenosine content in each captured sequence directly for accurate binding assessment. Adaptor and primer sequences used are detailed in the original publication.

### Single cells capture and library preparation (scRLP-seq)

Stable HEK293T cells were cultured in 10-cm dishes and induced with doxycycline (2 µg/mL, 24 h). Culture medium was carefully aspirated, and cells were washed twice with 3 mL of pre-warmed 1× PBS (37 °C) dispensed along the sidewall of the dish to avoid detachment. Cells were dissociated by incubation with 2 mL TrypLE Express (37 °C, Gibco) for 2 min. The reaction was neutralized by gently adding 5 mL of pre-warmed DMEM. Cells were pelleted (350 g, 3 min, RT), washed once with PBS and gently resuspended.

For viability assessment prior to ICELL8 loading, 1 mL of the resuspended cells was stained with a 1:1 mixture of Hoechst 33342 and propidium iodide (Thermo Fisher; 80 µL of each dye per 1 mL of cells). The suspension was mixed gently by inversion and incubated for 20 min at 37 °C protected from light. Stained cells were diluted with 1ml of ice-cold PBS, gently mixed, and pelleted (350 g, 3 min, 4 °C). The supernatant was carefully removed, and cells were resuspended in 1 mL of ice-cold PBS using a wide-bore pipette tip. Cell concentration was adjusted to 4 × 10⁴ cells/mL (1.4 cells/35 nL), suitable for dispensing into ICELL8 350v nanowell chips, with positive (K-562 RNA) and negative (PBS) controls included. Nanowells were imaged on the ICELL8 cx system, and single-cell candidates were first identified automatically with CellSelect v2.5 software based on Hoechst 33342–positive (blue, viable) nuclei and exclusion of propidium iodide–positive (red, dead) cells. All selections were then manually reviewed to ensure that only single, viable cells were retained for downstream processing.

Selected wells were subjected to first-strand cDNA synthesis using SMARTScribe™ reverse transcriptase and the SMART-Seq Pro oligo-dT primer with template switching oligonucleotide. Full-length cDNA was amplified by PCR (Terra™ Direct Polymerase, 13 cycles) on the ICELL8 cx thermal cycler. Amplified cDNAs were tagmented with bead-linked transposase (Illumina) and dual-indexed using SMART-Seq Pro Indexing Primer Plates. Libraries were extracted from the chip, purified with AMPure XP beads (Beckman), re-amplified, and purified again to yield sequencing-ready products.

Library quality was assessed using a High Sensitivity DNA5000 ScreenTape Assay (Agilent), and concentrations were quantified by qPCR of KAPA Library Quantification Kit (Roche). Pooled libraries were sequenced on an Illumina NovaSeq X Plus (2×150 bp), targeting 6-8 million paired reads per cell.

### Bulk RNA-seq and RNA editing detection

RNA-seq libraries were prepared using the NEBNext poly(A) mRNA Magnetic Isolation Module (E7490) following the manufacturer’s protocol. Libraries were sequenced on an Illumina NextSeq 2000/ NovaSeq X Plus platform with either single-end or paired-end reads (75 or 100-nt). Base calling and FASTQ conversion were performed using Illumina bcl2fastq (v2.20.0). Reads were trimmed with cutadapt (v4.9) and aligned to the Gencode v42 reference (GRCh38/hg38) using STAR (v2.7.9a) with the parameter --outFilterMultimapNmax 1 to retain only uniquely mapped reads. Redundant aligned reads were removed using Picard (v3.0.0). Gene expression level (TPM) was quantified using RSEM (v1.3.1). For **Figures 2B and S3C**, ERM-TadA-PABPC1 libraries were subsampled to a sequencing depth comparable to other reporters, whereas for **Figures 2C–2I and S3F-S3L** we applied deep sequencing (∼150 million reads) to maximize detection of RNAs associated with the ERM region.

RNA editing events were identified using JACUSA (v2.0.4), comparing doxycycline-induced samples against non-induced controls. The following parameters were applied: -a D (to filter potential false positives adjacent to INDEL positions, read start/end sites, and splice sites) - q2 30 (to exclude positions with base quality scores below 30). Events were retained if they passed all criteria: (1) absent from multiple non-induced controls, (2) only one SNV type (A-to-G or C-to-T) showing in this position, (3) supported by ≥3 A-to-G or C-to-U changes, and (4) contained ≥3 editing events with an editing ratio >0.01 per transcript.

Editing counts per million (ECPM) were calculated as:

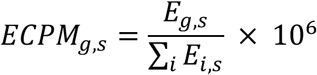

where 𝐸*_g,s_* is the total number of editing occurrences detected on all sequencing reads assigned to gene 𝑔 for sample 𝑠. Multiple edits on the same read were counted individually. The denominator ∑_..*i*_ 𝐸*_i,s_* represents the total number of editing occurrences across all genes in the sample 𝑠. Gene length was not normalized, as edit counts showed no correlation with transcript length (**Figure S1E**).

To identify the editing motif of APOBEC1-PABPC1-NES and TadA-PABPC1-NES, sequences spanning 10 nt upstream and downstream of each editing site were extracted and visualized by WebLogo3^48^. The 3′-UTR of human transcripts served as background for enrichment testing.

### Correlation analysis of compartment-specific RNA abundance with published datasets

To assess the concordance of compartment-specific RNA abundance between RLP measurements and published cytoplasmic/ER datasets, we compared ECPM values from corresponding RLP reporters with RNA-seq data from biochemical Fractionation-seq and FAPS studies. Genes with RPKM ≤ 1 were excluded to reduce the impact of low-abundance noise. Spearman correlation coefficients (ρ) were calculated between log₂(ECPM) and log₂(RPKM) across all genes. To examine expression-dependent trends, genes were further stratified into ten equal-width bins along the log₂(RPKM) axis, and the mean ± standard error of log₂(ECPM) was computed for each bin. Correlations were also assessed using these binned averages.

### Localization index calculation and temporal dynamics

To quantify RNA localization dynamics, we defined a localization index (LI) that compares the relative editing efficiency of each transcript between a target compartment and the cytoplasm. Editing efficiency was calculated as the ratio of total editing occurrences (edit counts, including multiple edits per read) to the total mapped read counts for each gene. For gene 𝑔,

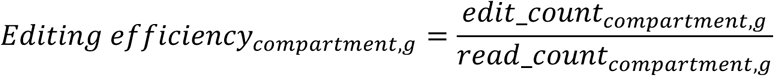

and similarly for the cytoplasm. The LI was then defined as:

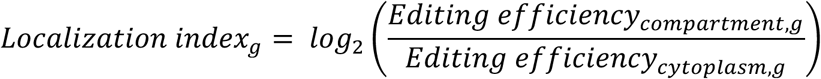

LI was calculated separately for ERM– and PM–targeted reporters, using matched cytoplasmic TadA-PABPC1-NES as the reference.

To characterize temporal patterns of RNA localization, the normalized LI was standardized using row-wise z-score normalization. Hierarchical clustering was then performed using average linkage (UPGMA algorithm) with correlation distance as the similarity metric. For visualization, the standardized scores were plotted as a heatmap with genes ordered according to the hierarchical clustering results.

### TranslationAI prediction of translation potential

To assess the translation potential of lncRNAs detected in cytoplasmic and ERM regions, we employed TranslationAI^32^, a deep learning-based prediction model specifically designed to identify functional translation initiation sites (https://www.biosino.org/TranslationAI/). Full-length sequences of the identified lncRNAs were submitted to the TranslationAI framework, which evaluates every nucleotide position for its probability of serving as a translation initiation site. For each lncRNA transcript, TranslationAI generated position-specific probability scores across the entire sequence. We extracted the maximum probability score from each transcript as its representative translation potential. Translation potential scores range from 0 to 1, with higher values indicating a greater probability that the RNA contains functional translation initiation sites that could lead to protein synthesis.

### Ribosome footprint data analysis

The ribosome profiling data analysis was performed using previously published ribosome footprint datasets from KEK293T cells (GEO accession: GSE97384)^49^. All analyses were conducted using the GRCh38 human genome assembly with GENCODE v42 annotations. The raw reads from three biological replicates of control sample were merged to increase coverage. Adapter sequences were trimmed using cutadapt (v4.9), and contaminating rRNA reads were removed by alignment to human rRNA sequences using Bowtie (v1.2.2). The remaining reads were aligned to the annotated transcriptome using STAR (v2.7.9a) with the parameter --outFilterMultimapNmax 1 to retain only uniquely mapped reads. Translation efficiency for each gene was calculated as the ratio of Ribo-seq read density to corresponding RNA-seq read density. Read densities were normalized as TPM using RSEM (v1.3.1), accounting for gene length and sequencing depth to enable direct comparison between conditions and transcript types.

### Gene functional analysis

Gene functional enrichment analysis was performed using the clusterProfiler (v4.0.5) package in R. Gene Ontology (GO) enrichment analysis was conducted for Biological Process (BP), Molecular Function (MF), and Cellular Component (CC) terms using the enrichGO function. The entire human genome gene set from org.Hs.eg.db was used as the background universe. Enrichment was calculated with the following parameters: Benjamini-Hochberg adjusted p-value cutoff of 0.05, and q-value cutoff of 0.05. To reduce redundancy in the enriched Gene Ontology (GO) terms, we applied the simplify function with a cutoff of 0.7 using the semantic similarity measure and selecting terms with the minimum adjusted p-value.

### Codon usage analysis

To investigate potential translational optimization patterns in compartment-specific transcripts, we analyzed the codon adaptation index (CAI) of genes localized to different cellular compartments. CAI values for each gene were obtained from the Codon Statistics Database (http://codonstatsdb.unr.edu). We extracted and compared codon usage patterns across gene sets corresponding to distinct subcellular localizations.

### Transmembrane domain and signal peptide prediction

To predict transmembrane domains and signal peptides in proteins whose corresponding RNAs are localized to different cellular compartments, we employed DeepTMHMM (v1.0)^50^, a deep learning model for transmembrane topology prediction and classification. Protein sequences for all genes in our dataset were retrieved from the UniProt database (release 2025_02). For each sequence, we performed predictions using DeepHMHMM with default parameters (confidence threshold = 0.7). Predictions were classified into four categories: (1) transmembrane proteins, (2) proteins with signal peptides only, (3) proteins with both features, and (4) proteins with neither feature.

### Conservation analysis

Transcript conservation was evaluated with PhastCons100 scores from 100-way vertebrate alignments (UCSC). For each transcript, the mean CDS conservation score was used in conservation analyses.

### UTR and protein lengths calculation

UTR and coding sequence (CDS) lengths were estimated from transcripts assembled with StringTie (v1.3.5) using RNA-seq data and the Gencode v42 (GRCh38/hg38) annotation as a reference. For genes with multiple expressed isoforms, lengths were calculated as the mean across all transcripts assembled for that gene. Protein length was defined as CDS length divided by three.

### scRLP-seq data analysis

Raw base call files were converted to FASTQ format using bcl-convert (v4.2.7) and demultiplexed with Cogent (v3.1), assigning reads to samples by library indexes and to single cells by ICELL8 well barcodes. Adapter sequences were trimmed with cutadapt (v4.9). Reads were aligned to the human reference genome (GRCh38/hg38, GENCODE v42) using STAR (v2.7.10b) in unique-alignment mode. PCR duplicates were removed with Picard (v2.27.1), and gene-level counts were quantified with featureCounts (v2.0.8) in unstranded paired-end mode (-s 0 -p –countReadPairs).

#### Cell quality control

Cells expressing fewer than 9,000 genes were excluded (genes were retained if supported by ≥5 unique reads in ≥2 cells). To ensure robust editing activity, cells were further filtered on A-to-G editing ratio and edited gene counts. Outliers were defined using the interquartile range (IQR) rule: for scRLP-NES, cells with values below Q1 – 1.5×IQR or above Q3 + 1.5×IQR; for scRLP-ERM, cells with <200 edited genes or >Q3 + 3×IQR were removed due to excessive heterogeneity. Because single cells were imaged, Hoechst⁺/PI-verified, and manually curated on the ICELL8 platform, and libraries were generated with SMART-seq2 (full-length, non-UMI), standard 10x-style filters based on UMI counts or mitochondrial fraction were not applied. Instead, QC relied on gene detection breadth (≥9,000 detected genes after deduplication), alignment quality, and editing-activity outlier removal as described above.

#### RNA editing detection

RNA editing events were identified with JACUSA (v2.0.4), -a D (to filter potential false positives adjacent to INDEL positions, read start/end sites, and splice sites), -q2 30 (to exclude positions with base quality scores below 30), and -P UNSTRANDED. Doxycycline-induced single-cell libraries were compared against non-induced bulk controls, and events overlapping control samples were removed. Editing events supported by fewer than three reads or in RNAs containing fewer than three edits were excluded.

#### Expression-based analysis

TPM–by–cell count matrices were processed with Seurat (v5.3.0, R 4.4.2). Data were log-normalized (*scale.factor=1e6*), 2,000 variable features were identified with the ‘*vst’* method, and data were scaled with ‘*ScaleData*’. Dimensionality reduction was performed by principal component analysis (PCA) conducted with ‘*RunPCA*’, and the top 20 principal components were selected for uniform manifold approximation and projection (UMAP) (*RunUMAP*, *n.neighbors=15*, *min.dist=0.1*) and for construction of the shared nearest neighbor graph. Clusters were identified with ‘*FindNeighbors*’ and ‘*FindClusters*’ (*resolution=0.3*). Cluster marker genes were defined with ‘*FindAllMarkers*’ (*min.pct=0.1, logfc.threshold=0.1*).

#### Editing-based analysis

Edit count–by–cell matrices were processed in parallel with Seurat. A Seurat object was created with *min.cells=3, min.features=0*, counts were log-normalized (*scale.factor=1e4*), and 2,000 variable editing sites were identified. PCA was performed, and the top 20 PCs were used for UMAP (*n.neighbors=15, min.dist=0.1, spread=0.5*) and clustering (*resolution=0.3*). Cluster-enriched editing sites were defined using ‘*FindAllMarkers*’ (*min.pct= 0.1, logfc.threshold=0.1*). For downstream analyses, the top 200 most variable cluster markers were used for Gene Ontology enrichment, ECPM distribution, and expression distribution analyses.

#### Cell-cycle analysis

Cell-cycle phases were assigned with Seurat’s CellCycleScoring using canonical S and G2/M markers (cc.genes). Phase distributions were tested by chi-square test, and S/G2M scores compared across phases by one-way ANOVA.

### Detection rate shuffling analysis

To evaluate whether scRLP-seq editing signals reflected true compartment-specific RNA localization rather than random labeling, we implemented a shuffling-based control. For each gene in the observed data, the detection rate was defined as the fraction of cells with ≥3 editing events, and genes were binned into 10% detection rate intervals. As a null model, we performed three independent shuffling experiments using the matched transcriptome (TPM) matrix. For each cell, we generated a synthetic read distribution in which total transcript abundance was proportional to TPM values, and then randomly reassigned the same number of editing events observed in the original editing matrix. Genes accumulating ≥3 simulated edits in at least one cell were retained for downstream analysis. For each shuffle, we computed gene detection rates across cells, binned them into 10% intervals as above, and recorded the number of labeled genes per bin. We summarized shuffled results by calculating the mean and standard deviation of gene counts per detection rate bin across the three shuffles.

Comparison of the observed and shuffled distributions provided a direct measure of how strongly scRLP-seq editing patterns deviated from random expectation. To formally evaluate this deviation, we applied a chi-square goodness-of-fit test, which compares the observed frequency distribution against the expected distribution from shuffling to determine whether they arise from the same underlying population.

### Detained intron ratio analysis

For each targeted intron in each single cell, we calculated the mean read coverage across the intron body, then normalized by the average of the mean coverages of the immediately upstream and downstream exons. This normalization controls for gene expression variability at the single-cell level and provides a normalized measure of intron presence relative to mature transcript signal. The same approach was applied to compute detained intron ratios in the APEX2-mediated proximity labeling dataset of nuclear domain–associated RNAs (GEO accession: GSE176439) ^45^.

### Published datasets used

Published datasets from HEK293T cells were used for comparison, including ER-enriched transcripts from APEX-seq^2^ (GEO accession: GSE116008), ER/cytosolic fractionation^17^ (GEO accession: GSE31539), fluorescence-activated particle sorting^5^ (GEO accession: GSE215770), and imaging-based MERFISH^30^, as well as RNA half-life measurements^51^ (GEO accession: GSE123165). SNHG family gene annotations were obtained from snoDB^52^.

### Statistics

Statistical comparisons were conducted with the Mann–Whitney U test unless noted otherwise. For enrichment analyses in pie charts, p-values were computed by a hypergeometric test with the background universe set at 20,000 human genes. All tests were two-sided unless otherwise specified.

## Acknowledgments

We are grateful to Gustavo Gutierrez-Cruz, Faiza Naz, Shamima Islam, and Dr. Stefania dell’Orso (NIAMS Genomics and Technology Section) for sequencing support. We also appreciate Davide Randazzo (NIAMS Light Imaging Section) for microscopy and imaging support, and Kevin Tinsley, Jeff Lay, and James Simone (NIAMS Flow Cytometry Section) for expert assistance with single live cell sorting. This research was supported by the Intramural Research Program of the National Institutes of Health (NIH). The contributions of the NIH authors are considered Works of the United States Government. The findings and conclusions presented in this paper are those of the authors and do not necessarily reflect the views of the NIH or the U.S. Department of Health and Human Services.

## Author Contributions

Conceptualization: M.H., X.F.; Methodology: X.F., and M.H.; Data analysis: X.F.; Experiments: X.F.; Writing and discussion: X.F., and M.H.; Funding acquisition: M.H..

## Declaration of Interests

None declared.

## Data availability

Raw bulk and single cell RNA-seq data from this study have been deposited in NCBI Gene Expression Omnibus (GEO) under accession code PRJNA1312488.

## Code availability

Source code and analysis scripts used in this study are publicly available at the GitHub repository: https://github.com/XJFan05/scRLP.

---

**Figure S1.**
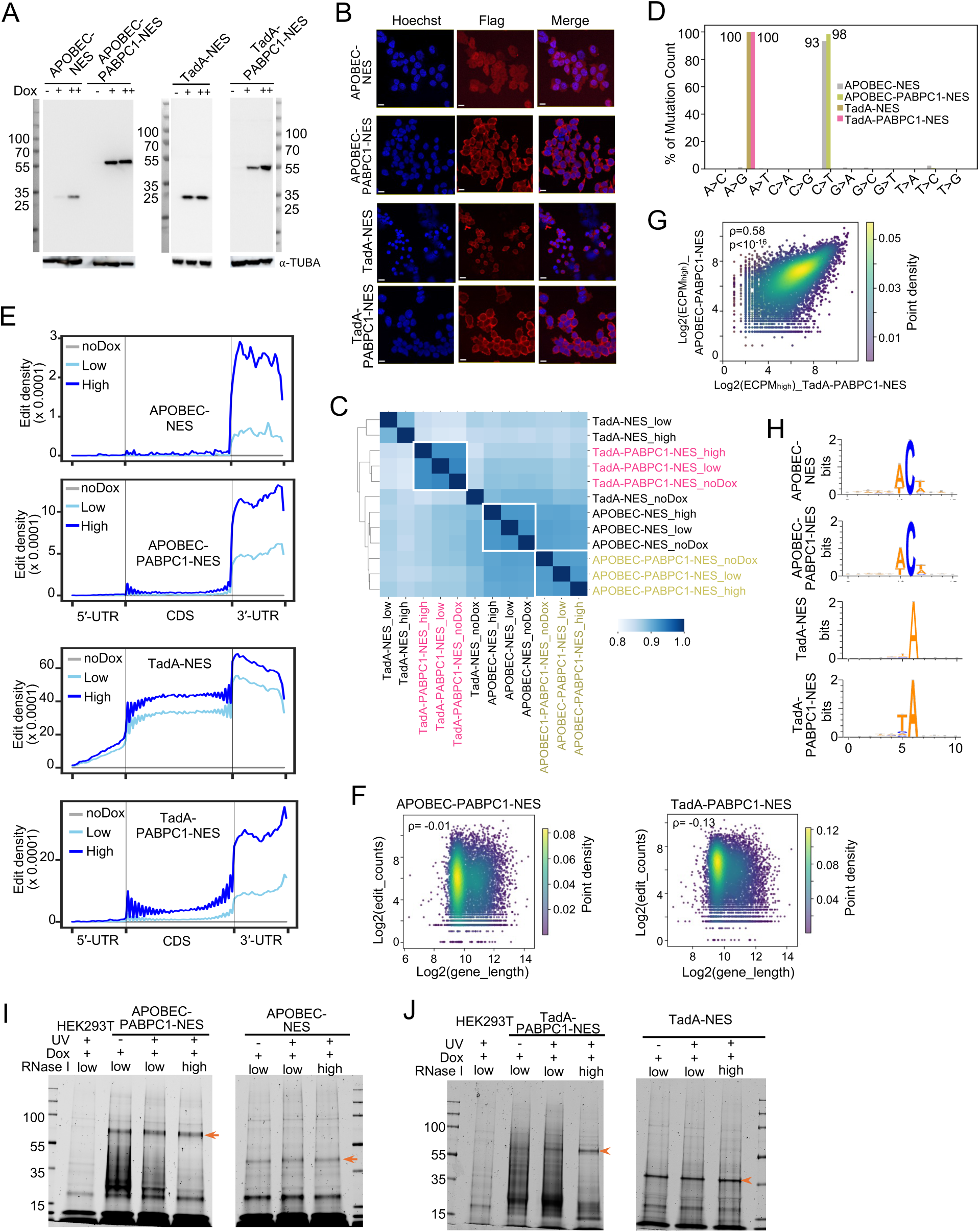
Characterization of RLP reporters in the cytoplasm. **A**, Western blot of APOBEC1-PABPC1-NES, TadA-PABPC1-NES, and control reporters lacking RBD of PABPC1 (APOBEC1-NES, TadA-NES) in stable cell lines 24 h after doxycycline induction, detected with anti-FLAG antibody. Cells were treated with no (−), low (+), or high (++) doxycycline. α-TUBA served as a loading control. Molecular weights (kDa) are indicated. **B**, Immunofluorescence (IF) images of the same reporters in panel **A**, collected 24 h after doxycycline induction. Nuclei were stained with Hoechst (blue) and reporter expression detected by FLAG immunostaining (red). Scale bars: 15 μm (APOBEC1-NES, APOBEC1-PABPC1-NES, and TadA-PABPC1-NES); 30 μm (TadA-NES). Fewer cells were observed in TadA-NES. **C**, Pairwise Pearson correlation heatmap of transcriptome profiles (log_10_ TPM) for each reporter under no, low, or high doxycycline induction. **D**, Single-nucleotide variant (SNV) distributions across 12 substitution types in cells expressing each reporter. **E**, Metagene plots of normalized editing density across the 5′ UTR, coding sequence (CDS), and 3′ UTR. Regions are scaled to relative length on the x-axis; normalized edit density is shown on the y-axis. Profiles are shown under no, low, and high doxycycline conditions. **F**, Scatter plots of log_2_ (gene length) *vs.* log_2_(edit counts) in cells expressing APOBEC1-PABPC1-NES (left) or TadA-PABPC1-NES (right). Spearman correlation coefficients (ρ) are indicated. **G**, Scatter plot of gene-level editing abundance (ECPM) between TadA-PABPC1-NES and APOBEC1-PABPC1-NES reporters under high doxycycline induction. Spearman correlation coefficients (ρ) and p value are indicated. **H**, Sequence logos showing nucleotide contexts surrounding editing sites for each reporter. **I-J**, Fluorescence imaging of PAR-CLIP gels for APOBEC1-PABPC1-NES and control APOBEC1-NES (panel **I**), or TadA-PABPC1-NES and control TadA-NES (panel **J**). Samples were prepared under no UV, low (0.015 U/µl), or high (0.15 U/µl) RNase I digestion. Untransfected controls were included. Orange arrows mark excised bands used for RNA-seq library preparation; corresponding gel regions from untransfected controls were processed in parallel.

**Figure S2.**
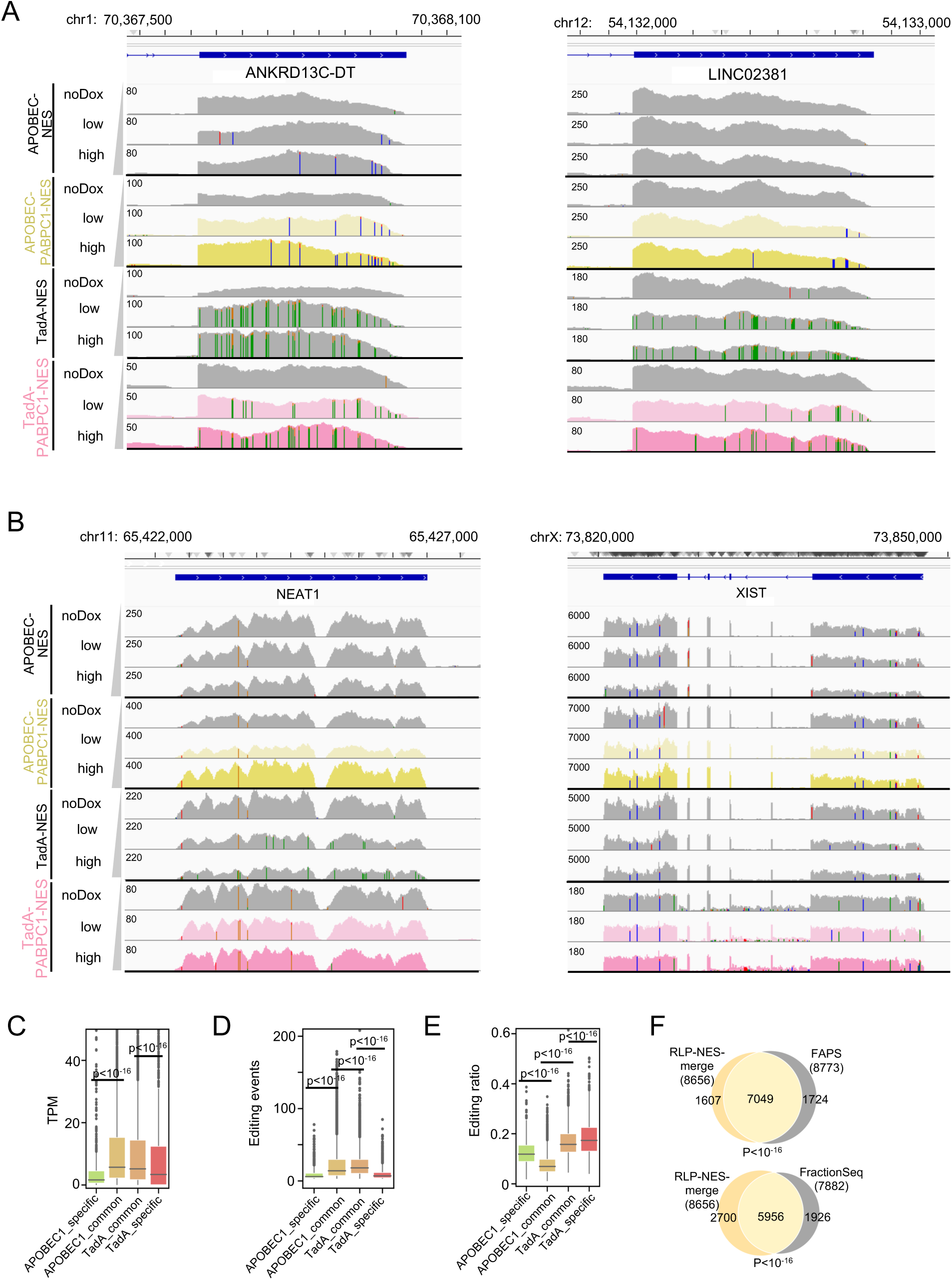
Characterization of cytoplasmic RNAs detected by RLP reporters. **A-B**, The editing profiles (RNA-seq coverage track) showing representative lncRNAs in cytoplasmic RLP datasets. Panel **A**, Representative edited lncRNAs ANKRD13C-DT and LINC02381, panel **B**, Representative non-edited lncRNAs NEAT1 and XIST. Read coverages are shown for APOBEC1-PABPC1-NES (yellow), TadA-PABPC1-NES (pink), and control (gray) reporters under no, low, or high doxycycline induction. Colored bars in coverage tracks indicate editing events (C-to-U for rAPOBEC1, A-to-G for TadA; SNV detection threshold = 0.05). Nucleotide colors: A = green, C = blue, G = orange, T = red (reversed on minus strand). **C-E**, Box plots comparing RNAs detected only by APOBEC1-PABPC1-NES (APOBEC1_specific), only by TadA-PABPC1-NES (TadA_specific), or by both reporters (APOBEC1_common, TadA_common). “APOBEC1_common” and “TadA_common” indicate shared RNAs, with features taken from the APOBEC1-PABPC1-NES and TadA-PABPC1-NES datasets, respectively. Panel **C**, Transcript abundance (TPM). Panel **D**, Number of editing events. Panel **E**, Editing ratio. P values are indicated. **F**, Overlap of cytoplasmic RNAs detected by both RLP reporters with those identified in published cytosolic fractionation datasets. P values are indicated.

**Figure S3.**
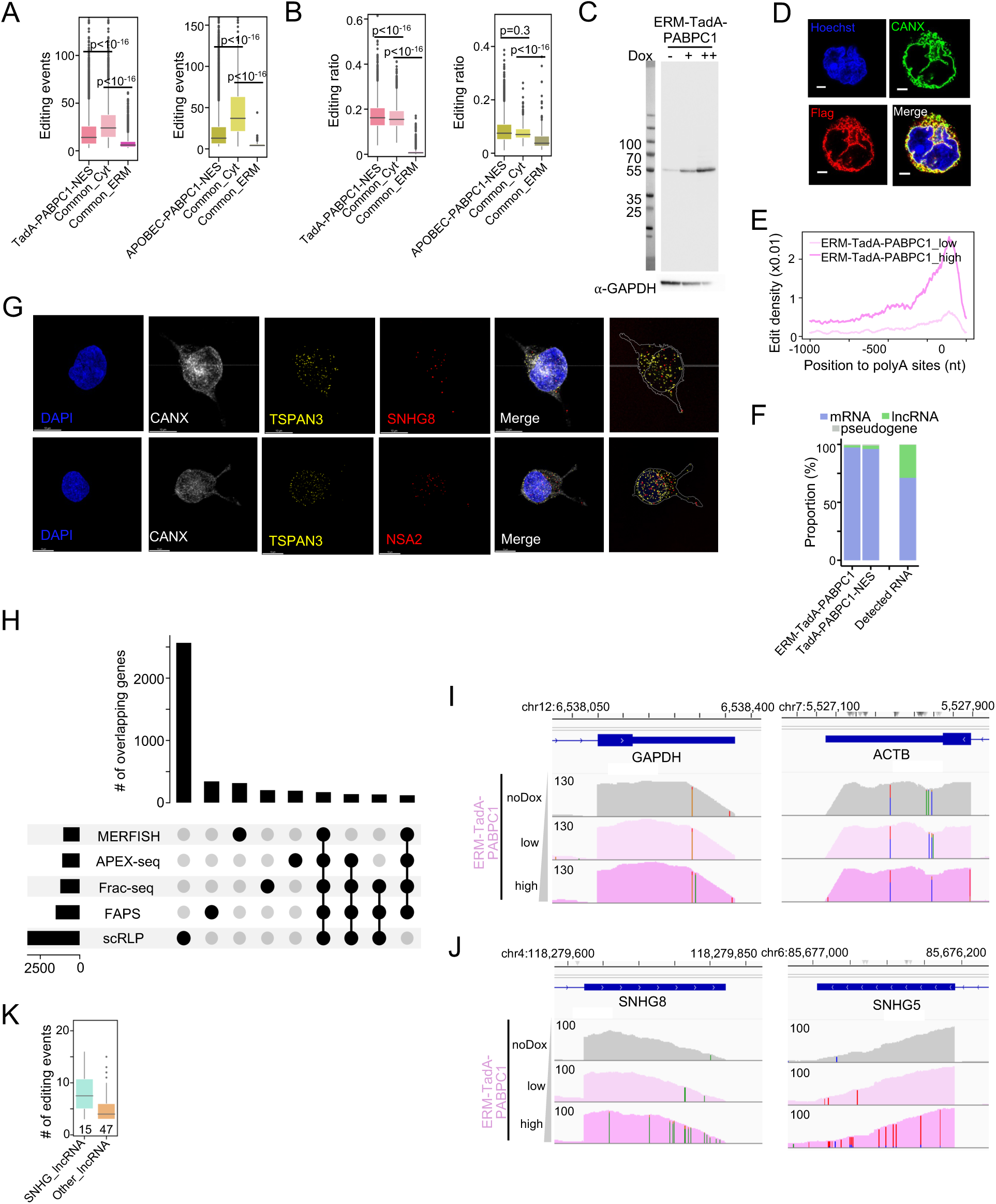
Characterization of the RLP reporter in the endoplasmic reticulum membrane (ERM). **A-B**, Distribution of (panel **A**) editing events and (panel **B**) editing ratios for three RNA groups: RNAs detected only by TadA-PABPC1-NES or APOBEC1-PABPC1-NES in the cytoplasm, RNAs shared between cytoplasm and ERM with values from TadA-PABPC1-NES or APOBEC1-PABPC1-NES (Common_Cyt), and RNAs shared between cytoplasm and ERM with values from ERM-TadA-PABPC1 or ERM-APOBEC1-PABPC1 (Common_ERM). P values are indicated. **C**, Western blot of ERM-TadA-PABPC1 expression in stable cell line 24 h after doxycycline induction, detected with anti-FLAG antibody. Cells were treated with no (−), low (+), or high (++) doxycycline. α-GAPDH served as a loading control; Molecular weights (kDa) are indicated on the left. **D**, IF images of ERM-TadA-PABPC1 reporter 24 h after induction. Nuclei stained with Hoechst (blue), ER with CANX (green), and reporter with FLAG immunostaining (red). Scale bars, 2 μm. **E**, Metagene analysis of editing density relative to poly(A) sites. **F**, Composition of detected ERM-associated RNAs compared with cytoplasm RNAs and total RNA-seq. Bars show the proportions of edited mRNAs (blue), lncRNAs (green), and pseudogenes (gray) in ERM-TadA-PABPC1, TadA-PABPC1-NES, and total detected RNAs by RNA-seq. **G**, Representative multiplex FISH-IF validation of ERM-associated RNAs from z-stack images. DAPI (blue), CANX (gray), ERM control RNA TSPAN3 (yellow), and test ERM RNAs (red; SNHG8 and NSA2). Rightmost panels show nuclear and ER masks (gray) overlaid with RNA puncta (yellow/red). Scale bar, 10 µm. See **Supplementary Videos 4–5**. **H**, Overlap of ERM-associated transcripts identified by RLP, FAPS, APEX-seq, Fractionation-seq (Frac-seq), and MERFISH. The top nine intersections by gene count are shown. **I-J**, The editing profiles (RNA-seq coverage track) of panel **I**, GAPDH and ACTB and panel **J**, representative ERM-associated SNHG family RNAs SNHG8 and SNHG5, from ERM-TadA-PABPC1 reporter under no, low, or high doxycycline induction. Colored bars indicate editing events (SNV detection threshold = 0.05). **K**, The number of editing events detected in SNHG family lncRNAs compared with other ERM-associated lncRNAs.

**Figure S4.**
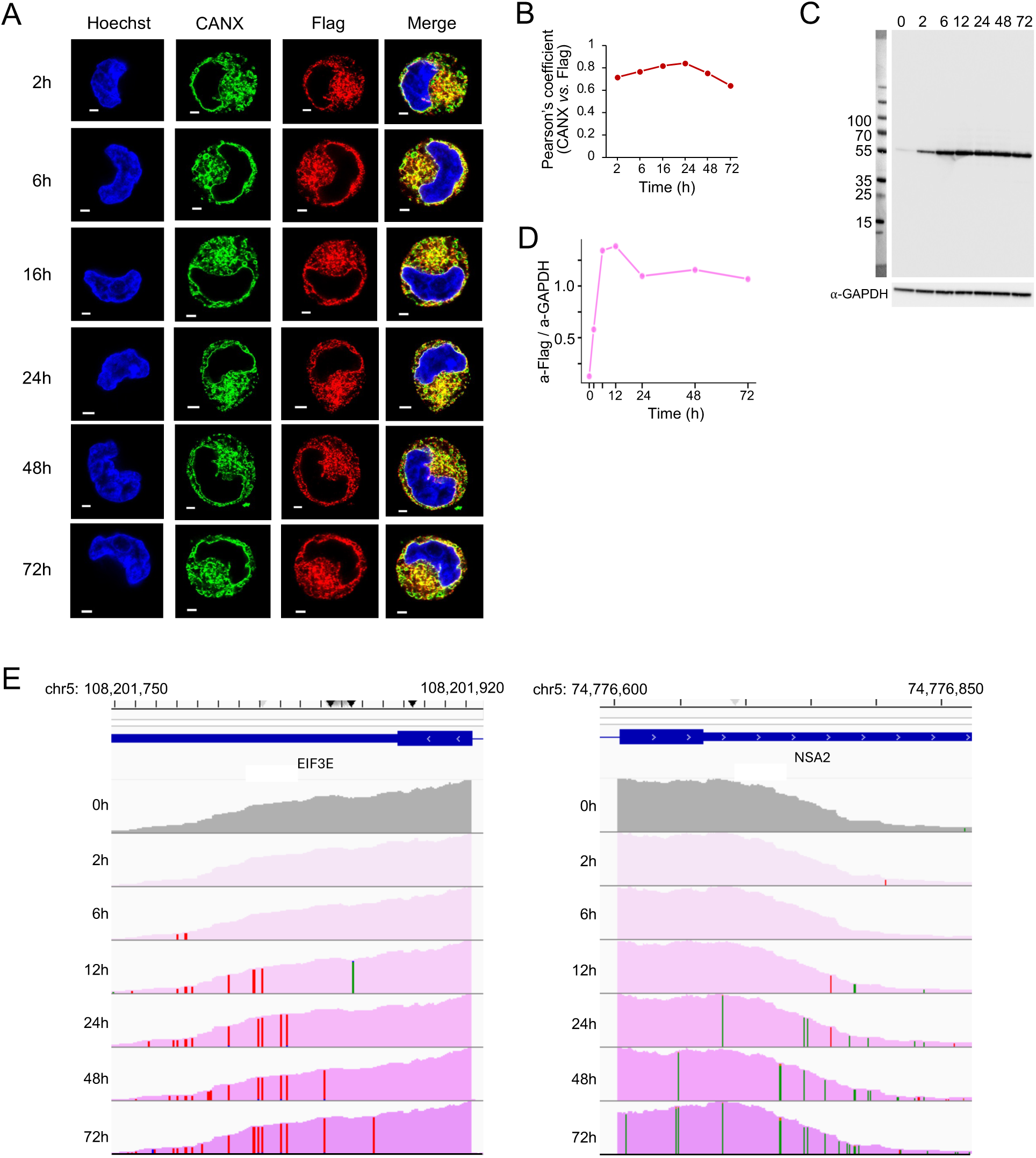
Time-course analysis of the RLP reporters at the ERM. **A**, IF images of ERM-TadA-PABPC1 from 2-72 h after doxycycline induction. Nuclei were stained with Hoechst (blue), ER with CANX (green), and reporter with FLAG immunostaining (red). Scale bars, 2 μm. **B**, Pearson’s correlation coefficients between CANX (green) and FLAG (red) signals in 3D Z-stacks acquired on the Leica TCS SP8. N ≥ 8 cells per time point. **C**, Western blot of ERM-TadA-PABPC1 expression from 0-72h after induction (0 h, uninduced control), detected with anti-FLAG antibody. α-GAPDH served as a loading control; molecular weights (kDa) are indicated. **D**, Quantification of FLAG signal normalized to GAPDH (α-FLAG/α-GAPDH) at the indicated times, corresponding to panel **C**. **E**, The editing profiles (RNA-seq coverage track) of representative ERM-associated cluster 2 RNAs (EIF3E and NSA2). Colored bars indicate editing events (A-to-G for TadA; SNV detection threshold = 0.02).

**Figure S5.**
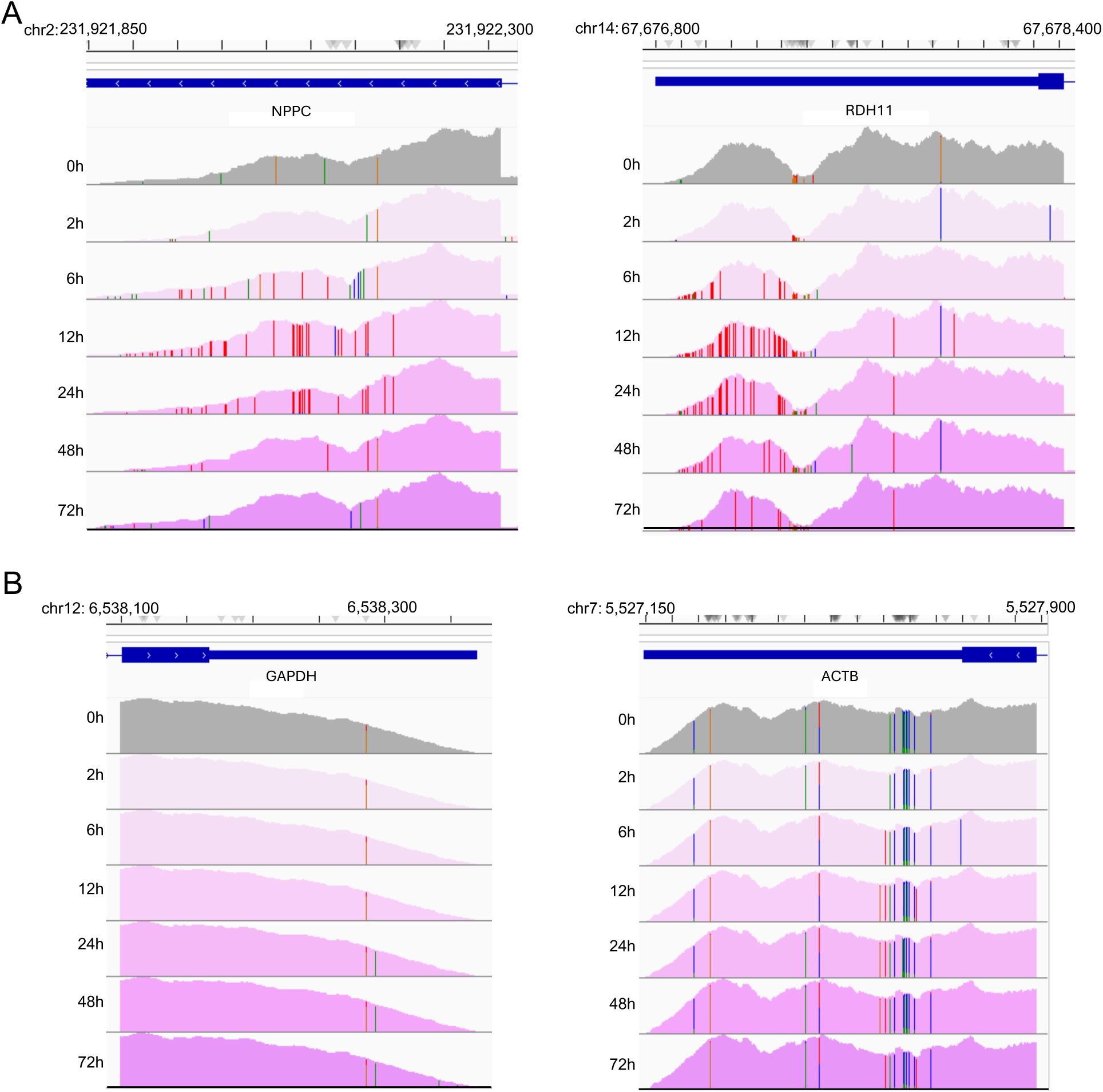
The editing profiles of representative RNAs. **A**, ERM-associated cluster 4 RNAs NPPC and RDH11. **B**, Cytoplasmic RNAs GAPDH and ACTB. For both panels, read coverages are shown for ERM-TadA-PABPC1 at 0 h (gray) and 2–72 h (purple). Colored bars indicate editing events (A-to-G for TadA; SNV detection threshold = 0.02).

**Figure S6.**
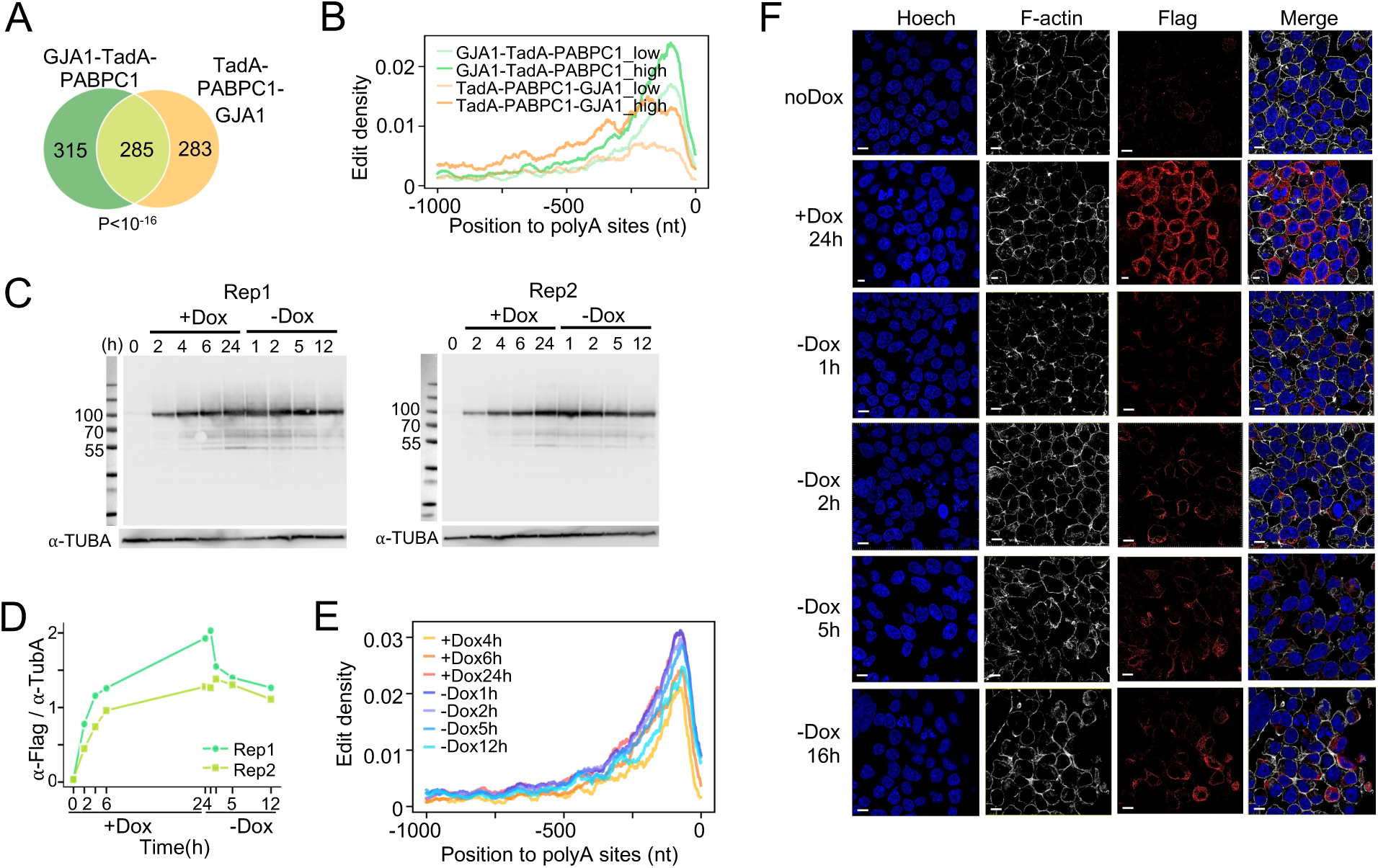
Characterization of RLP reporter at the plasma membrane (PM). **A**, Overlap of RNAs identified by GJA1-TadA-PABPC1 and TadA-PABPC1-GJA1 reporters. P values are indicated. **B**, Metagene profiles of editing density relative to poly(A) sites for GJA1-TadA-PABPC1 and TadA-PABPC1-GJA1 reporters. **C**, Western blot of GJA1-TadA-PABPC1 reporter across the dual-phase experiment in two biological replicates. The reporter was detected with anti-FLAG antibody. α-TUBA served as a loading control; molecular weights (kDa) are indicated. **D**, Quantification of reporter expression by FLAG signal normalized to α-TUBA (α-FLAG/α-TUBA) at the indicated times, corresponding to panel **C**. **E**, Metagene profiles of editing density relative to poly(A) sites during the dual-phase experiment. **F**, IF images of GJA1-TadA-PABPC1 reporter in the dual-phase experiment. Nuclei are stained with Hoechst (blue), F-actin with phalloidin (gray), and reporter with FLAG immunostaining (red). A high-resolution single-cell image acquired 1 h after doxycycline removal is shown in **Supplementary Video 6.** Scale bars, 10 μm.

**Figure S7.**
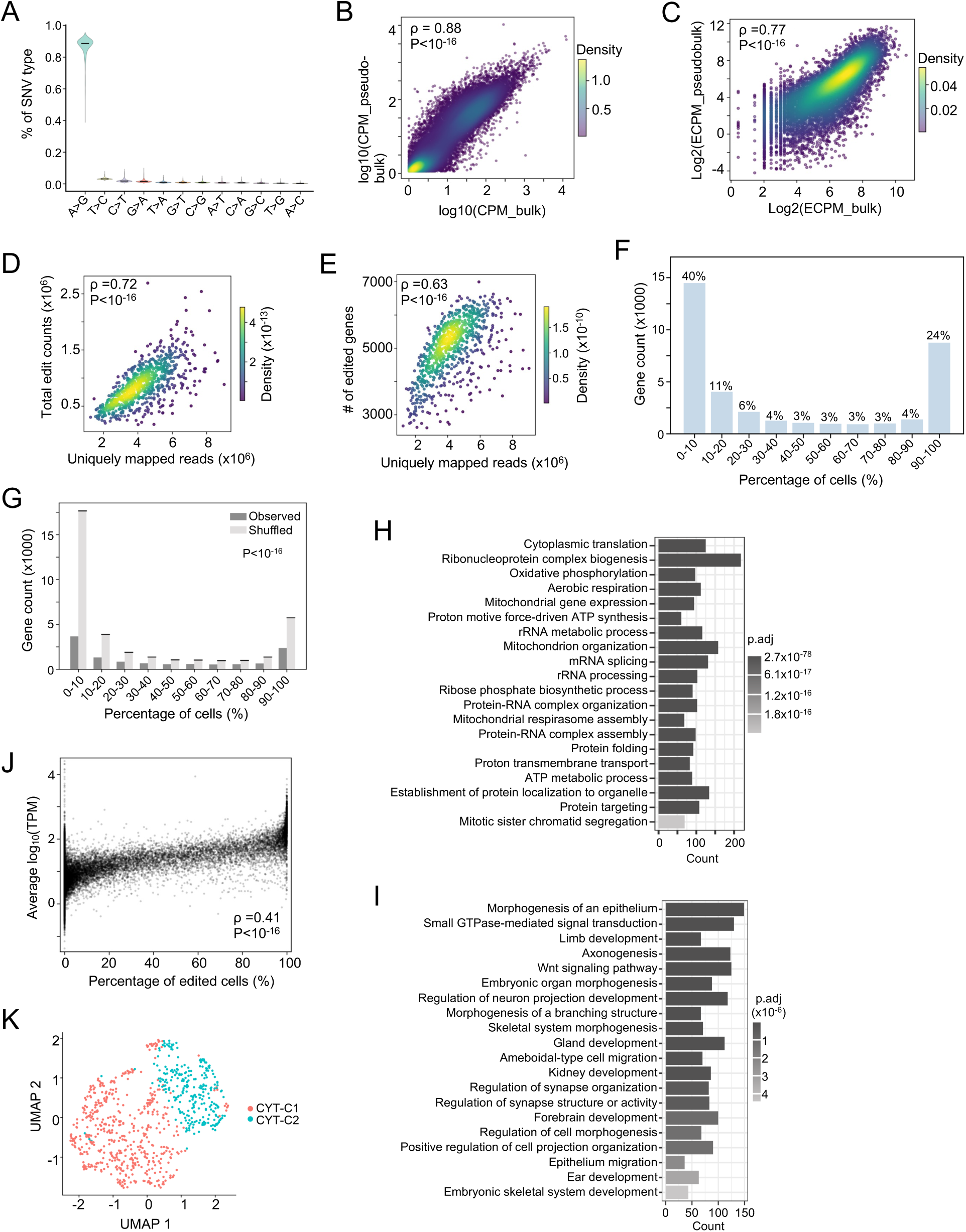
Characterization of cytoplasmic RNAs detected by scRLP-seq. **A**, SNV distributions across 12 substitution types among single cells expressing TadA-PABPC1-NES. **B**, Correlation of expression between bulk and pseudobulk scRLP-seq (log_10_ CPM). Spearman ρ and p value are indicated. **C**, Correlation of editing level between bulk and pseudobulk scRLP-seq data (log_2_ ECPM). Spearman ρ and p value are indicated. **D-E**, Scatter plots of uniquely mapped reads *vs.* edit counts (panel **D**) or edited genes (panel **E**) per cell. **F**, Distribution of all detected genes by scRLP-seq (y-axis) binned by detection frequency across cells (x-axis). Percentages relative to the total number of detected genes are shown above. **G**, Genes were binned by detection rate and compared with three independent shuffling-based controls. Bars show observed data (gray) and shuffled controls (light gray, mean ± SD). **H-I**, GO Biological Process enrichment of RNAs detected in >90% (panel **H**) or <10% (panel **I**) of cells. **J**, Relationship between RNA abundance and the detection frequency across cells. **K**, UMAP of single cells based on editing profiles.

**Figure S8.**
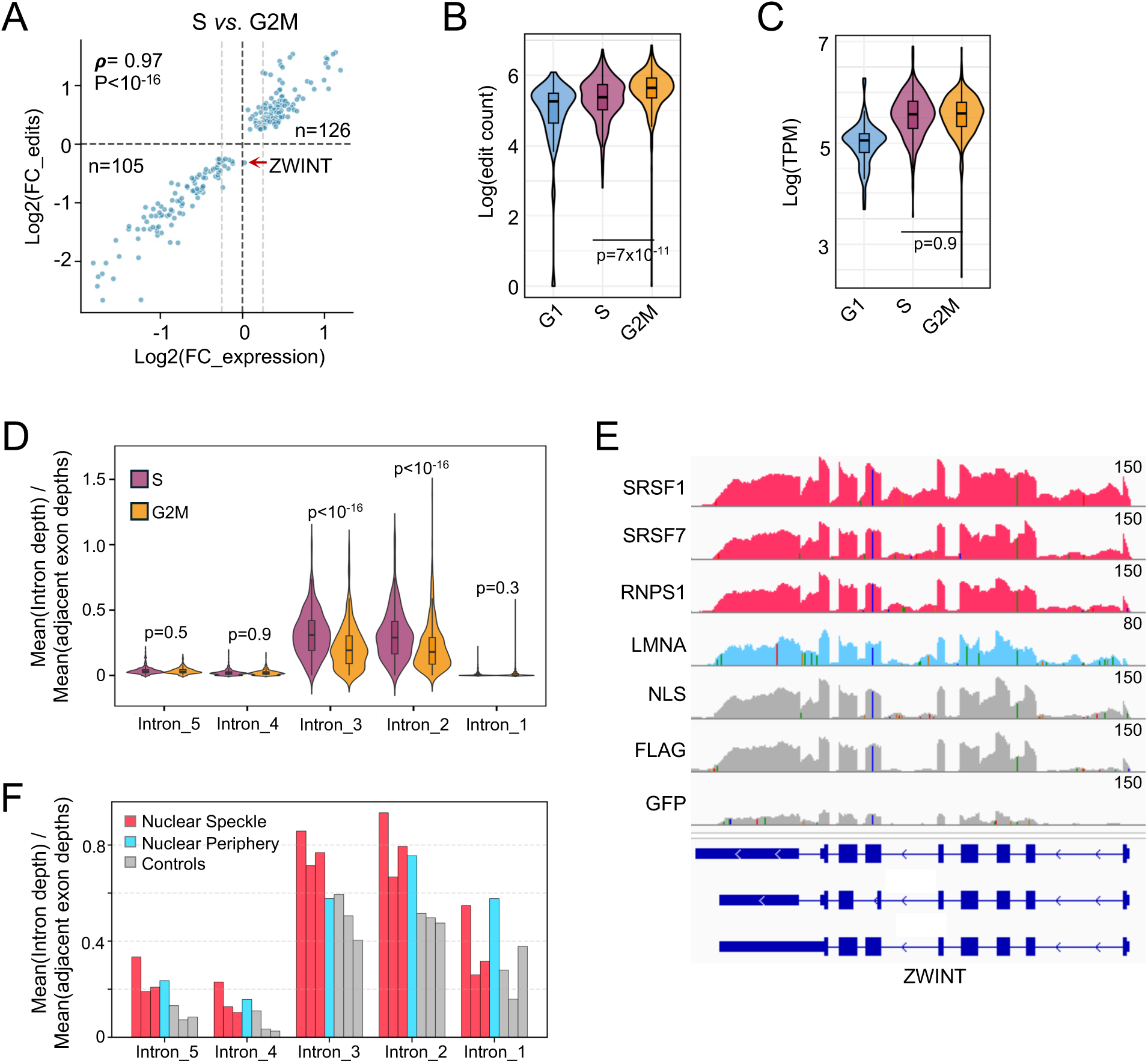
Redistribution of ZWINT mRNA during the S–G2/M transition. **A**, Scatter plot of log_2_ fold change (FC) of expression (x-axis) *vs.* editing (y-axis) for genes differing between S and G2/M phases (adjusted p < 0.05; |log₂ FC| ≥ 0.25). Dashed lines indicate zero and ±0.25 thresholds. Spearman ρ and p value are shown. Quadrant counts indicate gene numbers. ZWINT is highlighted (red arrow). **B-C**, The distributions of ZWINT editing (panel **B**) and expression (panel **C**) across single cells in G1, S, and G2/M phases. **D**, Distribution of intron ratios for constitutive introns of ZWINT across single cells measured by scRLP-seq. **E**, Genome tracks showing raw RNA-seq reads from the APEX2-nuclear marker and control samples mapped to the ZWINT locus. **F**, Intron ratio of ZWINT calculated from panel **E**. Nuclear speckle markers: SRSF1, SRSF7, and RNPS1; nuclear periphery marker: LMNA; controls: NLS, FLAG, and GFP.

**Figure S9.**
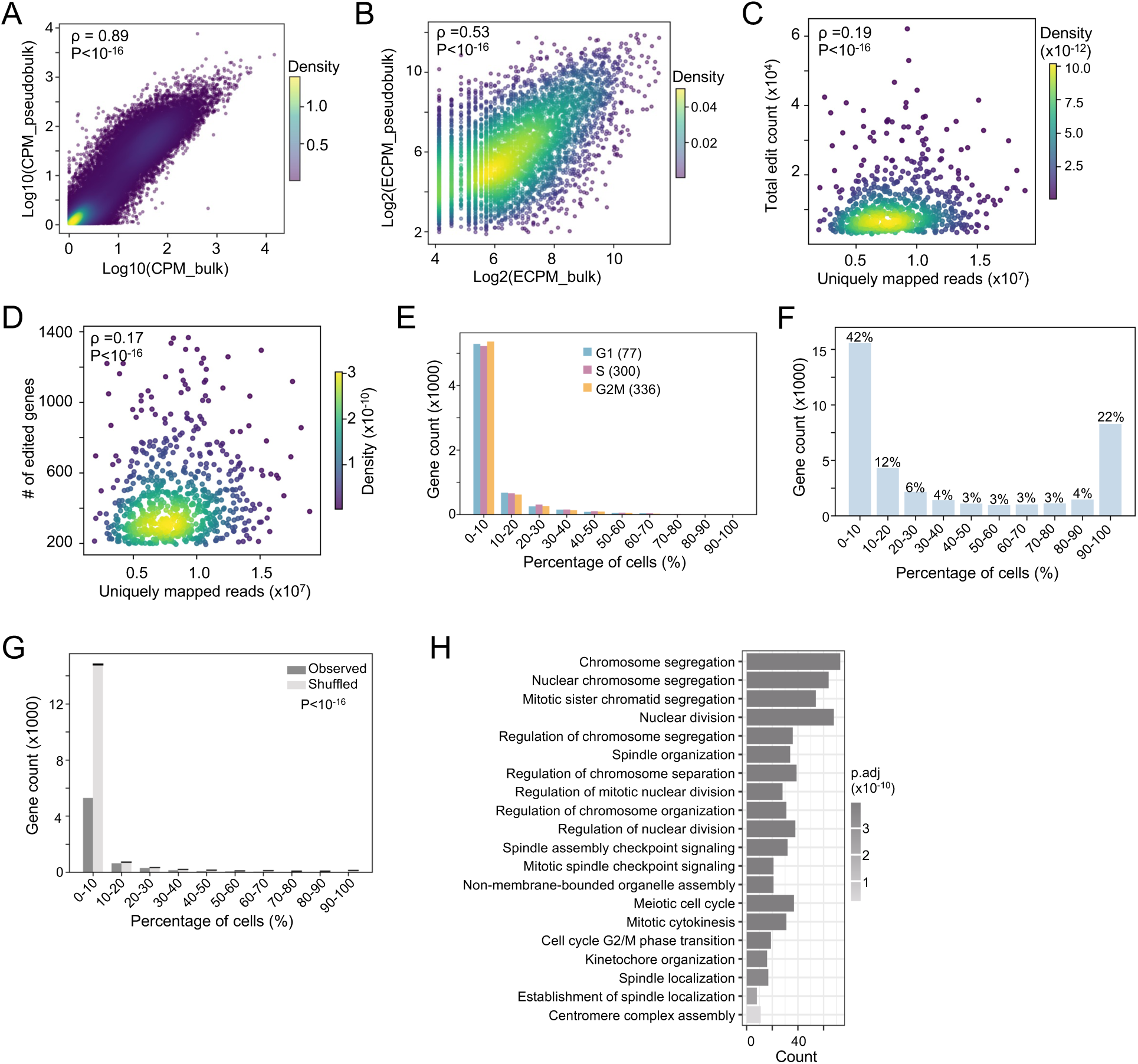
Characterization of ERM-associated RNAs detected by scRLP-seq. **A-B**, Correlation analysis of expression (log_10_ CPM, panel **A**) or editing (log_2_ ECPM, panel **B**) between bulk and pseudobulk scRLP-seq data. Spearman ρ and p value are indicated. **C-D**, Scatter plots of uniquely mapped reads *vs.* edit counts (panel **C**) or edited genes (panel **D**) per cell. **E**, Distribution of edited gene counts (y-axis) by detection frequency across cells (x-axis) for G1, S, and G2/M phases. **F**, Distribution of all genes detected by scRLP-seq (y-axis) binned by detection frequency across cells (x-axis). Percentages relative to the total number of detected genes are shown above. **G**, Genes were binned by detection rate (fraction of cells with ≥3 editing events) in scRLP-seq data and compared with three independent shuffling simulations based on TPM-derived read distributions. Bars show the number of edited genes per bin for the observed data (gray) and the shuffled controls (light gray, mean ± SD). **H**, GO Biological Process enrichment of the top 200 most variable genes from the expression-based UMAP in Figure 7D.

## Notes

### Competing Interest Statement

The authors have declared no competing interest.

## References

1. Banani, S.F., Lee, H.O., Hyman, A.A., and Rosen, M.K. (2017). Biomolecular condensates: organizers of cellular biochemistry. Nat Rev Mol Cell Biol 18, 285–298. 10.1038/nrm.2017.7.

2. Fazal, F.M., Han, S., Parker, K.R., Kaewsapsak, P., Xu, J., Boettiger, A.N., Chang, H.Y., and Ting, A.Y. (2019). Atlas of Subcellular RNA Localization Revealed by APEX-Seq. Cell 178, 473–490 e426. 10.1016/j.cell.2019.05.027.

3. Hristov, P., and Flynn, R.A. (2024). Imaging glycosylated RNAs at the subcellular scale. Nat Biotechnol 42, 574–575. 10.1038/s41587-023-02021-1.

4. Perr, J., Langen, A., Almahayni, K., Nestola, G., Chai, P., Lebedenko, C.G., Volk, R.F., Detres, D., Caldwell, R.M., Spiekermann, M., et al. (2025). RNA-binding proteins and glycoRNAs form domains on the cell surface for cell-penetrating peptide entry. Cell 188, 1878–1895 e1825. 10.1016/j.cell.2025.01.040.

5. Horste, E.L., Fansler, M.M., Cai, T., Chen, X., Mitschka, S., Zhen, G., Lee, F.C.Y., Ule, J., and Mayr, C. (2023). Subcytoplasmic location of translation controls protein output. Mol Cell 83, 4509–4523 e4511. 10.1016/j.molcel.2023.11.025.

6. Holt, C.E., Martin, K.C., and Schuman, E.M. (2019). Local translation in neurons: visualization and function. Nat Struct Mol Biol 26, 557–566. 10.1038/s41594-019-0263-5.

7. Holt, C.E., and Bullock, S.L. (2009). Subcellular mRNA localization in animal cells and why it matters. Science 326, 1212–1216. 10.1126/science.1176488.

8. Mili, S., Moissoglu, K., and Macara, I.G. (2008). Genome-wide screen reveals APC-associated RNAs enriched in cell protrusions. Nature 453, 115–119. 10.1038/nature06888.

9. Mardakheh, F.K., Paul, A., Kumper, S., Sadok, A., Paterson, H., McCarthy, A., Yuan, Y., and Marshall, C.J. (2015). Global Analysis of mRNA, Translation, and Protein Localization: Local Translation Is a Key Regulator of Cell Protrusions. Dev Cell 35, 344–357. 10.1016/j.devcel.2015.10.005.

10. Martin, K.C., and Ephrussi, A. (2009). mRNA localization: gene expression in the spatial dimension. Cell 136, 719–730. 10.1016/j.cell.2009.01.044.

11. Gasparski, A.N., Moissoglu, K., Pallikkuth, S., Meydan, S., Guydosh, N.R., and Mili, S. (2023). mRNA location and translation rate determine protein targeting to dual destinations. Mol Cell 83, 2726–2738 e2729. 10.1016/j.molcel.2023.06.036.

12. Moissoglu, K., Yasuda, K., Wang, T., Chrisafis, G., and Mili, S. (2019). Translational regulation of protrusion-localized RNAs involves silencing and clustering after transport. Elife 8. 10.7554/eLife.44752.

13. Cabili, M.N., Dunagin, M.C., McClanahan, P.D., Biaesch, A., Padovan-Merhar, O., Regev, A., Rinn, J.L., and Raj, A. (2015). Localization and abundance analysis of human lncRNAs at single-cell and single-molecule resolution. Genome Biol 16, 20. 10.1186/s13059-015-0586-4.

14. Ren, J., Zhou, H., Zeng, H., Wang, C.K., Huang, J., Qiu, X., Sui, X., Li, Q., Wu, X., Lin, Z., et al. (2023). Spatiotemporally resolved transcriptomics reveals the subcellular RNA kinetic landscape. Nat Methods 20, 695–705. 10.1038/s41592-023-01829-8.

15. Samacoits, A., Chouaib, R., Safieddine, A., Traboulsi, A.M., Ouyang, W., Zimmer, C., Peter, M., Bertrand, E., Walter, T., and Mueller, F. (2018). A computational framework to study sub-cellular RNA localization. Nat Commun 9, 4584. 10.1038/s41467-018-06868-w.

16. Cajigas, I.J., Tushev, G., Will, T.J., tom Dieck, S., Fuerst, N., and Schuman, E.M. (2012). The local transcriptome in the synaptic neuropil revealed by deep sequencing and high-resolution imaging. Neuron 74, 453–466. 10.1016/j.neuron.2012.02.036.

17. Reid, D.W., and Nicchitta, C.V. (2012). Primary role for endoplasmic reticulum-bound ribosomes in cellular translation identified by ribosome profiling. J Biol Chem 287, 5518–5527. 10.1074/jbc.M111.312280.

18. Engel, K.L., Lo, H.G., Goering, R., Li, Y., Spitale, R.C., and Taliaferro, J.M. (2022). Analysis of subcellular transcriptomes by RNA proximity labeling with Halo-seq. Nucleic Acids Res 50, e24. 10.1093/nar/gkab1185.

19. Benhalevy, D., Anastasakis, D.G., and Hafner, M. (2018). Proximity-CLIP provides a snapshot of protein-occupied RNA elements in subcellular compartments. Nat Methods 15, 1074–1082. 10.1038/s41592-018-0220-y.

20. Chen, K.H., Boettiger, A.N., Moffitt, J.R., Wang, S., and Zhuang, X. (2015). Spatially resolved, highly multiplexed RNA profiling in single cells. Science 348, aaa6090. 10.1126/science.aaa6090.

21. Eng, C.L., Lawson, M., Zhu, Q., Dries, R., Koulena, N., Takei, Y., Yun, J., Cronin, C., Karp, C., Yuan, G.C., and Cai, L. (2019). Transcriptome-scale super-resolved imaging in tissues by RNA seqFISH+. Nature 568, 235–239. 10.1038/s41586-019-1049-y.

22. McMahon, A.C., Rahman, R., Jin, H., Shen, J.L., Fieldsend, A., Luo, W., and Rosbash, M. (2016). TRIBE: Hijacking an RNA-Editing Enzyme to Identify Cell-Specific Targets of RNA-Binding Proteins. Cell 165, 742–753. 10.1016/j.cell.2016.03.007.

23. Navaratnam, N., Morrison, J.R., Bhattacharya, S., Patel, D., Funahashi, T., Giannoni, F., Teng, B.B., Davidson, N.O., and Scott, J. (1993). The p27 catalytic subunit of the apolipoprotein B mRNA editing enzyme is a cytidine deaminase. J Biol Chem 268, 20709–20712.

24. Tegowski, M., Flamand, M.N., and Meyer, K.D. (2022). scDART-seq reveals distinct m(6)A signatures and mRNA methylation heterogeneity in single cells. Mol Cell 82, 868–878 e810. 10.1016/j.molcel.2021.12.038.

25. Brannan, K.W., Chaim, I.A., Marina, R.J., Yee, B.A., Kofman, E.R., Lorenz, D.A., Jagannatha, P., Dong, K.D., Madrigal, A.A., Underwood, J.G., and Yeo, G.W. (2021). Robust single-cell discovery of RNA targets of RNA-binding proteins and ribosomes. Nat Methods 18, 507–519. 10.1038/s41592-021-01128-0.

26. Medina-Munoz, H.C., Kofman, E., Jagannatha, P., Boyle, E.A., Yu, T., Jones, K.L., Mueller, J.R., Lykins, G.D., Doudna, A.T., Park, S.S., et al. (2024). Expanded palette of RNA base editors for comprehensive RBP-RNA interactome studies. Nat Commun 15, 875. 10.1038/s41467-024-45009-4.

27. Xiao, Y.L., Liu, S., Ge, R., Wu, Y., He, C., Chen, M., and Tang, W. (2023). Transcriptome-wide profiling and quantification of N(6)-methyladenosine by enzyme-assisted adenosine deamination. Nat Biotechnol 41, 993–1003. 10.1038/s41587-022-01587-6.

28. Deo, R.C., Bonanno, J.B., Sonenberg, N., and Burley, S.K. (1999). Recognition of polyadenylate RNA by the poly(A)-binding protein. Cell 98, 835–845. 10.1016/s0092-8674(00)81517-2.

29. Hafner, M., Landthaler, M., Burger, L., Khorshid, M., Hausser, J., Berninger, P., Rothballer, A., Ascano, M., Jr., Jungkamp, A.C., Munschauer, M., et al. (2010). Transcriptome-wide identification of RNA-binding protein and microRNA target sites by PAR-CLIP. Cell 141, 129–141. 10.1016/j.cell.2010.03.009.

30. Xia, C., Fan, J., Emanuel, G., Hao, J., and Zhuang, X. (2019). Spatial transcriptome profiling by MERFISH reveals subcellular RNA compartmentalization and cell cycle-dependent gene expression. Proc Natl Acad Sci U S A 116, 19490–19499. 10.1073/pnas.1912459116.

31. Dominguez, D., Freese, P., Alexis, M.S., Su, A., Hochman, M., Palden, T., Bazile, C., Lambert, N.J., Van Nostrand, E.L., Pratt, G.A., et al. (2018). Sequence, Structure, and Context Preferences of Human RNA Binding Proteins. Mol Cell 70, 854–867 e859. 10.1016/j.molcel.2018.05.001.

32. Fan, X., Chang, T., Chen, C., Hafner, M., and Wang, Z. (2025). Analysis of RNA translation with a deep learning architecture provides new insight into translation control. Nucleic Acids Res 53. 10.1093/nar/gkaf277.

33. Monziani, A., and Ulitsky, I. (2023). Noncoding snoRNA host genes are a distinct subclass of long noncoding RNAs. Trends Genet 39, 908–923. 10.1016/j.tig.2023.09.001.

34. Dermit, M., Dodel, M., Lee, F.C.Y., Azman, M.S., Schwenzer, H., Jones, J.L., Blagden, S.P., Ule, J., and Mardakheh, F.K. (2020). Subcellular mRNA Localization Regulates Ribosome Biogenesis in Migrating Cells. Dev Cell 55, 298–313 e210. 10.1016/j.devcel.2020.10.006.

35. Ribeiro-Rodrigues, T.M., Martins-Marques, T., Morel, S., Kwak, B.R., and Girao, H. (2017). Role of connexin 43 in different forms of intercellular communication - gap junctions, extracellular vesicles and tunnelling nanotubes. J Cell Sci 130, 3619–3630. 10.1242/jcs.200667.

36. Beardslee, M.A., Laing, J.G., Beyer, E.C., and Saffitz, J.E. (1998). Rapid turnover of connexin43 in the adult rat heart. Circ Res 83, 629–635. 10.1161/01.res.83.6.629.

37. Komar, A.A., Samatova, E., and Rodnina, M.V. (2024). Translation Rates and Protein Folding. J Mol Biol 436, 168384. 10.1016/j.jmb.2023.168384.

38. Cheng, L.C., Zheng, D., Zhang, Q., Guvenek, A., Cheng, H., and Tian, B. (2021). Alternative 3’ UTRs play a widespread role in translation-independent mRNA association with the endoplasmic reticulum. Cell Rep 36, 109407. 10.1016/j.celrep.2021.109407.

39. Weinberg, D.E., Shah, P., Eichhorn, S.W., Hussmann, J.A., Plotkin, J.B., and Bartel, D.P. (2016). Improved Ribosome-Footprint and mRNA Measurements Provide Insights into Dynamics and Regulation of Yeast Translation. Cell Rep 14, 1787–1799. 10.1016/j.celrep.2016.01.043.

40. Picelli, S., Faridani, O.R., Bjorklund, A.K., Winberg, G., Sagasser, S., and Sandberg, R. (2014). Full-length RNA-seq from single cells using Smart-seq2. Nat Protoc 9, 171–181. 10.1038/nprot.2014.006.

41. Wang, H., Hu, X., Ding, X., Dou, Z., Yang, Z., Shaw, A.W., Teng, M., Cleveland, D.W., Goldberg, M.L., Niu, L., and Yao, X. (2004). Human Zwint-1 specifies localization of Zeste White 10 to kinetochores and is essential for mitotic checkpoint signaling. J Biol Chem 279, 54590–54598. 10.1074/jbc.M407588200.

42. Braun, C.J., Stanciu, M., Boutz, P.L., Patterson, J.C., Calligaris, D., Higuchi, F., Neupane, R., Fenoglio, S., Cahill, D.P., Wakimoto, H., et al. (2017). Coordinated Splicing of Regulatory Detained Introns within Oncogenic Transcripts Creates an Exploitable Vulnerability in Malignant Glioma. Cancer Cell 32, 411–426 e411. 10.1016/j.ccell.2017.08.018.

43. Boutz, P.L., Bhutkar, A., and Sharp, P.A. (2015). Detained introns are a novel, widespread class of post-transcriptionally spliced introns. Genes Dev 29, 63–80. 10.1101/gad.247361.114.

44. Dominguez, D., Tsai, Y.H., Weatheritt, R., Wang, Y., Blencowe, B.J., and Wang, Z. (2016). An extensive program of periodic alternative splicing linked to cell cycle progression. Elife 5. 10.7554/eLife.10288.

45. Barutcu, A.R., Wu, M., Braunschweig, U., Dyakov, B.J.A., Luo, Z., Turner, K.M., Durbic, T., Lin, Z.Y., Weatheritt, R.J., Maass, P.G., et al. (2022). Systematic mapping of nuclear domain-associated transcripts reveals speckles and lamina as hubs of functionally distinct retained introns. Mol Cell 82, 1035–1052 e1039. 10.1016/j.molcel.2021.12.010.

46. Lam, S.S., Martell, J.D., Kamer, K.J., Deerinck, T.J., Ellisman, M.H., Mootha, V.K., and Ting, A.Y. (2015). Directed evolution of APEX2 for electron microscopy and proximity labeling. Nat Methods 12, 51–54. 10.1038/nmeth.3179.

47. Anastasakis, D.G., Jacob, A., Konstantinidou, P., Meguro, K., Claypool, D., Cekan, P., Haase, A.D., and Hafner, M. (2021). A non-radioactive, improved PAR-CLIP and small RNA cDNA library preparation protocol. Nucleic Acids Res 49, e45. 10.1093/nar/gkab011.

48. Crooks, G.E., Hon, G., Chandonia, J.M., and Brenner, S.E. (2004). WebLogo: a sequence logo generator. Genome Res 14, 1188–1190. 10.1101/gr.849004.

49. Park, Y., Reyna-Neyra, A., Philippe, L., and Thoreen, C.C. (2017). mTORC1 Balances Cellular Amino Acid Supply with Demand for Protein Synthesis through Post-transcriptional Control of ATF4. Cell Rep 19, 1083–1090. 10.1016/j.celrep.2017.04.042.

50. Jeppe Hallgren, K.D.T., Mads D. Pedersen, José Juan Almagro Armenteros, Paolo Marcatili, Henrik Nielsen, Anders Krogh and Ole Winther (2022). DeepTMHMM predicts alpha and beta transmembrane proteins using deep neural networks. bioRxiv. 10.1101/2022.04.08.487609.

51. Narula, A., Ellis, J., Taliaferro, J.M., and Rissland, O.S. (2019). Coding regions affect mRNA stability in human cells. RNA 25, 1751–1764. 10.1261/rna.073239.119.

52. Bergeron, D., Paraqindes, H., Fafard-Couture, E., Deschamps-Francoeur, G., Faucher-Giguere, L., Bouchard-Bourelle, P., Abou Elela, S., Catez, F., Marcel, V., and Scott, M.S. (2023). snoDB 2.0: an enhanced interactive database, specializing in human snoRNAs. Nucleic Acids Res 51, D291–D296. 10.1093/nar/gkac835.

